# Comparative metagenomic study unveils bacterial communities and their putative involvement in ecological success of two pine-feeding *Ips* beetle holobionts

**DOI:** 10.1101/2024.02.23.581803

**Authors:** Arunabha Khara, Amrita Chakraborty, Roman Modlinger, Amit Roy

**Author notes:** Corresponding Author: Amit Roy, Amrita Chakraborty.

## Abstract

Climate change has recently boosted the severity and frequency of the pine bark beetle attacks. The bacterial community associated with these beetles acts as “hidden players”, enhancing their ability to infest and thrive on defence-rich pine trees. There is limited understanding of the environmental acquisition of these hidden players and their life stage-specific association with different pine-feeding bark beetles. There is inadequate knowledge on novel bacterial introduction to pine trees after the beetle infestation. Hence, we conducted the first comparative bacterial metabarcoding study comprehensively revealing the bacterial communities in the pine trees before and after beetle feeding and in different life stages of two dominant pine-feeding bark beetles, namely *Ips sexdentatus* and *Ips acuminatus*. We also evaluated the bacterial association between wild and lab-bred beetles to measure the deviation due to inhabiting a controlled environment. Significant differences in bacterial amplicon sequence variance (ASVs) abundance existed among different life stages within and between the pine beetles. Such observations endorsed that the bark beetle life stage shaped bacterial assemblage. Furthermore, lab-bred and wild-collected adult beetles had distinct bacterial assemblages, implying that the breeding environment induced crucial changes. Alteration of pine wood bacteriome after beetle feeding is an intriguing observation in the present study, which demands further investigation. We validated the relative abundances of selected bacterial taxa estimated by metagenomic sequencing with quantitative PCR. Functional predictions revealed that these bacterial genera might execute conserved functions, aiding the ecological success of these beetles. Nevertheless, these findings shed new insights into bacterial associations and their putative metabolic roles in pine beetles under the influence of various drivers such as environment, host, and life stages and provide the foundation for future downstream functional investigations.

**Importance:** The current understanding of bark beetle as holobiont is restricted. Most studies lack information on microbial community assembly in bark beetle microhabitats. No data comprehensively reveals the influence of lab breeding on pine beetle microbial associations. It is unknown if there is any adaptive convergence in beetle microbial assemblage due to feeding on the same host. Such information is essential to developing a bark beetle management strategy to restore forests from beetle-mediated damage. Our study shows that lab-breeding considerably influences beetle bacterial community assembly. We documented that beetle feeding alters bacteriome at the microhabitat level, and the beetle life stage shapes the bacterial associations. Nevertheless, our study revisited the bark beetle symbiosis under the influence of different drivers and revealed intriguing insight into bacterial community assembly, facilitating future functional studies.

## INTRODUCTION

Bark beetles (*Coleoptera: Curculionidae: Scolytinae*) are economically important forest pest that causes large-scale forest damage across Europe (1). The outbreaks of pine bark beetles such as *Ips sexdentatus* (Börner, 1776) and *Ips acuminatus* (Gyllenhal, 1827) heavily impact forestry economics, affecting the forest-dependent sector and international wood markets (2). These outbreaks reduce the forest tree lifespan, decrease carbon uptake, and alter microclimatic situations in the forest, along with recreational and aesthetic values (3, 4). According to recent reports, particularly in the Czech Republic, the severity of pinewood damage by bark beetles increased by over 80,000 m³ in 2019 from about 10,000 m³ in 2009 (5). Moreover, climate changes such as increased air temperature, altered precipitation patterns, and increased frequency of drought and heat events have critically stimulated the outbreaks of bark beetles by altering the tree defence physiology (6, 7). Precisely, drought affects host tree fitness and can stimulate bark-beetle populations to overcome the epidemic threshold and cause an outbreak (8). For instance, high spring temperature stimulates *I. acuminatus* to infest weakened and vigorous trees (9). Subsequently, the increasing infestations by *I. acuminatus* have listed the species among the most-aggressive bark beetle species within Europe (5, 10–12). Climate change has also stimulated the importance of *I. sexdentatus* as a forest pest with considerable dispersal competency (13).

Although *I. sexdentatus* and *I. acuminatus* share several ecological characteristics, including host, *I. sexdentatus* infests the lower part of the bole, whereas *I. acuminatus* attacks the higher bark canopy (top branches) (14). Nonetheless, both *Ips* species must overcome the robust pine defence system to thrive. The robust pine defence system has physical and chemical barriers against biotic stress, including bark beetles (15). Physical barriers include tissues having lignin and suberin polymers that offer protection against degradation, penetration and ingestion, while the chemical barriers involve phenolic compounds and terpenoids (monoterpenes, diterpenes, and sesquiterpenes) with entomotoxic competency. A high concentration of monoterpenes shows ovicidal, repellent, adulticidal, and larvicidal effects against bark beetles (16). Therefore, the defence barrier in healthy pine trees is a formidable challenge for bark beetles. Alternatively, microbial assemblage in bark beetles promotes the competency of bark beetles to exhaust conifer defence. For instance, bacterial genera such as *Pseudomonas*, *Serratia*, and *Rahnella* associated with *Dendroctonus valens* (LeConte, 1860) metabolized monoterpenes (17, 18). Similarly, *Erwinia typography* isolated from *Ips typographus* (Linnaeus, 1758) can tolerate a high concentration of myrcene-a plant defensive monoterpene (19). Furthermore, mountain pine beetle (*Dendroctonus ponderosae* Hopkins) has been reported to have bacterial mutualists containing genes (i.e., dit genes-diterpene degrading genes) associated with terpene degradation (20).

Apart from overcoming the tree defence, a significant challenge for bark beetles is nutrient acquisition, as they primarily thrive on phloem tissues with limited nutrition and complex carbohydrates. Subsequently, the gut microbiome comes to the rescue, helping the beetles to digest and metabolize complex polysaccharides (i.e., lignin from conifers) (21) and nutrient acquirement (22, 23). Beetle-associated microbiome extends their symbiosis with the host by producing pheromones, thus facilitating chemical communications (24). However, such information on microbial contribution to bark beetle survival is limited to only a few beetle genera, such as the red turpentine and mountain pine beetle (20, 25). Nevertheless, our understanding of bark beetles as holobionts is restricted. Few studies have evaluated microbial acquisition and succession in bark beetles (26–29). In addition, limited studies in pine bark beetles have focused on understanding the effect of environment and metamorphosis that can cause comprehensive changes in the beetle microbiome, resulting in distinct bacterial communities. Moreover, microbial community assembly at the level of bark beetle microhabitat is still lacking. There is limited information on the influence of the host plant microbiome in shaping the beetle bacterial community. No literature highlights the impact of beetle feeding on the pine-associated microbiome. Subsequently, it is also crucial to assess the microbial assembly in wild and lab-bred beetles to evaluate the influence of the bark beetle breeding facility.

We hypothesized that metamorphosis, microhabitat, and lab-breeding might shape the bacterial community structure in the *Ips* pine beetles similar to other insects (27–32). Hence, we conducted the first comparative metabarcoding study on two pine-feeding beetles (*I. sexdentatus*, *I. acuminatus*) to evaluate our hypothesis and respond to the following fundamental questions regarding bark beetle symbiosis: (1) What are the bacterial communities associated with the two pine-feeding *Ips* beetles across life-stages? (2) What influence does the host microbiome have in shaping the bacterial community assemblage or *vice versa*? (3) How does lab-breeding impact pine beetle bacterial community assemblage?

## MATERIAL AND METHODS

### Bark beetle collection and breeding for life-stage specific sample collection

Adult bark beetles (Coleoptera: Curculionidae: Scolytinae) were collected from infested logs (dbh ∼20 cm) obtained from different beetle colonies in the Rouchovany forest area (49.0704° N, 16.1076° E) in the Czech Republic in 2020. The two bark beetle species (*I.sexdentatus* and *I.acuminatus*) were identified based on the work of Nunberg (33) and Pfeffer (34, 35). The infested logs were brought to the debarking room directly from the forest, where the adult beetles were collected using surface disinfected tweezers and bred to F2 generation in laboratory conditions (temp around 25°C in the day and 19°C at night, RH 60%) on fresh pine logs collected from the same forest area. Over 50 samples from each life stage (larva, pupa, adult) were collected from multiple infested logs in 50 ml plastic conical tubes, snap-frozen with liquid nitrogen, and kept at −80℃ for DNA extraction. The gallery wood was collected from the same F2 generation infested logs using surface sterilized blades and kept in plastic tubes with RNAlater solution at −80℃ for future use. All the beetle and wood samples were collected at least 10 cm below the edges of the logs to minimize contamination. Uninfected, fresh woods with no beetle infestation were taken as control wood (unfed fresh phloem) and were kept in plastic conical tubes with RNAlater solution at −80℃.

### Wild beetle collection

Similarly, the infested logs, collected from the same Rouchovany forest area in 2021, were directly brought to the debarking room, where adult *I. sexdentatus* and *I. acuminatus* wild beetles were collected using surface sterilized tweezers in 50 ml plastic conical tubes, snap-frozen with liquid nitrogen and kept at −80℃ for DNA extraction. The gallery wood samples were collected from the same infested logs in RNAlater solution and stored at −80℃. Similarly, unfed wood samples (fresh phloem tissue) were collected from logs from the same locality without any beetle infestation. Beetle bacteriome variability of individual colonies was not explored in the current study due to the mixing of beetle samples from different colonies. However, such a sampling method can reduce the random variability from the heterologous sampling material. The sample details are provided in **Table 1**.

**Table 1:**
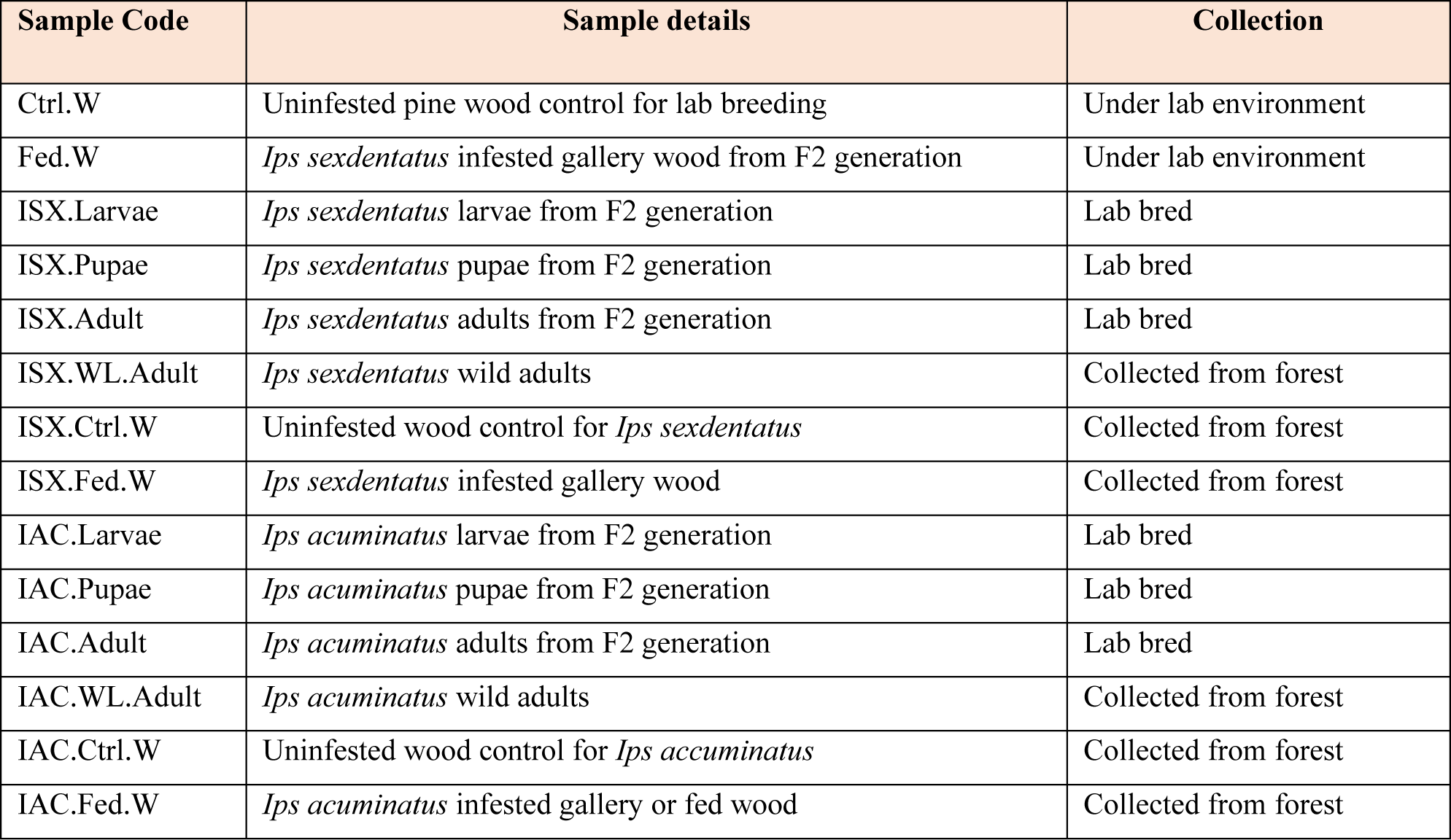
Sample description.

### DNA extraction

Developmental stage-specific lab-bred and wild-collected samples were randomly selected and disinfected with 70% ethanol. The ethanol was removed by rinsing it with sterile water. Any sample with apparent infection was discarded. Due to size differences, the sampling amount per replicate for both species differed. One individual per replicate was used for *I. sexdentatus*, whereas, for *I. acuminatus,* 4 larvae/replicate, 2 pupae/replicate, and 5 adults/replicate were used. Total DNA from beetles was extracted using the MACHEREY-NAGEL NucleoSpin Soil DNA kit with modifications in the manufacturer’s protocol. Similarly, wood microbial DNA was extracted using QIAGEN DNeasy Plant Mini Kit following the manufacturer’s protocol with modifications. The samples were homogenized under liquid nitrogen, and the lysis was performed for 1 minute. The extracted DNA quantity and quality were accessed using a Qubit 2.0 Fluorometer (Thermo Scientific) and 1% agarose gel electrophoresis. High-quality samples (5 biological replicates from each life stage for both species and 4 biological replicates for each wood sample) and two negative extraction controls were selected for 16S rRNA metabarcoding at Novogene Company, China.

### 16S amplicon sequencing

16S amplicon sequencing was executed at Novogene, China, using a pre-optimized protocol. Precisely, 1ng/µl template DNA, bacterial 16S rRNA gene primers (341F-806R) (36) containing unique barcodes, and Phusion High-Fidelity PCR Master Mix (New England Biolabs) were used to set up PCR reactions. Subsequently, a PCR reaction mixture without template DNA was used as a negative control. PCR products were visualized in 2% agarose gel electrophoresis, and bands ranging between 450-470 bp were excised from the gel using a QIAGEN Gel extraction kit. NEBNext Ultra DNA Library Pre-Kit for Illumina was used to create sequencing libraries followed by index code ligation. Sequencing libraries were quantitively and qualitatively analyzed using Qubit 2.0 Fluorometer (Thermo Fisher Scientific) and Agilent Bioanalyser 2100 system. An Illumina Novaseq6000 platform was used to obtain 250 bp paired-end reads from the sequenced libraries.

### Bacteriome Data Analysis

#### Data processing and species annotation

The bioinformatic data analysis was performed using a standardized pipeline in QIIME2 (version 2022.2) (37) as described in our earlier studies (27). The barcodes and primer sequences were removed, and the sample sequences were merged using FLASH (V1.2.11, http://ccb.jhu.edu/software/FLASH/) to generate raw Illumina pair-end reads (38) and then checked for high-quality reads using fastp software. VSEARCH software (39) was used to identify and remove chimeric sequences to facilitate downstream bioinformatic analyses. The amplicon sequence variant (ASV) (40) abundance table was obtained using the DADA2 module (41). The ASV abundance table or the feature table is a matrix of samples, and the feature (ASV) abundance represents the number of times each feature/ASV was observed in each sample. Sequences that have an abundance lesser than five were discarded. Furthermore, the ASVs were compared with the bacterial SILVA database (version 138) (42) to obtain individual ASVs with respective species annotation in QIIME2 (version 2022.2) (37).

#### Alpha diversity

Alpha diversity indices like bacterial community richness (Chao1), evenness (Pielou) (43) and bacterial diversity (Shannon, Simpson) (43) were used to analyze bacterial community structure. Kruskal-Wallis-pairwise-group test was performed to test the significance level within the samples. In addition, QIIME2 (version 2022.2) was used to estimate Good’s coverage (sequence depth) (44) and observed species from all stage-specific and wood samples that were represented by R software (Version 2.15.3; R Core Team, 2013, Vienna, Austria) (45).

#### Beta diversity

The bacterial diversity variation between different life stages and wood samples for both *Ips* species was estimated using the UniFrac distance metric (46) determined in QIIME2 (version 2022.2). Moreover, unweighted UniFrac distance measurement was used for non-metric multi-dimensional scaling (NMDS) analysis illustrated by R software (47). The functions ADONIS (analysis and partitioning sum of squares using dissimilarities) and ANOSIM (analysis of similarities) (48, 49) were used to determine significant differences in life-stage specific and wood bacteriome in QIIME2 (version 2022.2) (Supplementary Table 1 and Supplementary Table 2). ADONIS is a non-parametric multivariate variance test analysis that utilizes a distance metric (46) to determine significant differences in the bacterial community among the sample groups (50). However, ANOSIM analysis utilizes the same distance metric to estimate if the variation among different sample groups is larger than within the sample group (51). Furthermore, a t-test was used to determine significantly abundant bacterial species (p-value < 0.05) in the sample groups (52). Similarly, Metastats analysis using false discovery rate (FDR) and multiple hypothesis tests for sparsely-sampled features revealed that the intra-group differed significantly on abundant bacterial species (53). LEfSe (linear discriminant analysis effect size) analysis was performed to obtain significant biomarkers that can help to distinguish two samples in an experimental condition (54). These biologically consistent, statistically significant biomarkers derived from LEfSe can disclose metagenomic attributes (taxa/metabolites/genes) to distinguish between the two samples.

Finally, PICRUSt2 software (Phylogenetic Investigation of Communities by Reconstruction of Unobserved Stats 2) (55) and the bacterial ASVs tree with relative gene information were used to extrapolate its probable functional profile. The known and unknown bacterial groups obtained from the sequences that matched the SILVA database were compared with KEGG (Kyoto Encyclopaedia of Genes and Genomes) (56) and COG (Clusters of Orthologous Genes) (57) databases to generate heatmaps containing the probable functions of the bacterial groups. Furthermore, the significant abundance of putative functional genes based on the KEGG database was predicted. The relative abundance of Enzyme Commission (EC) between two life stages at p<0.05 with the KEGG Enzyme database was used to develop comparative heatmaps to visualize their difference in abundance between the life stages of both pine beetles. The heatmaps are categorized according to three fundamental functions (amino acid biosynthesis, cell wall degradation, vitamins, and co-factors biosynthesis) essential for beetle development and survival.

### Quantitative PCR assay

The relative abundance of selected bacterial taxa was estimated using quantitative PCR assay and correlated with the metagenomic sequencing results. Six individuals for each life stage (larvae/pupae/adults) were pooled per replicate, and four biological replicates were prepared. However, due to the limited availability of *I. acuminatus* samples, only four replicates of adult beetles were prepared. Due to the same reason, the life stage comparisons for *I. acuminatus* were not performed using qPCR. Tubulin beta-1 chain (β*-Tubulin*) was used as reference genes for *I. sexdentatus* (58), while ribosomal protein (*RPL*7) and the elongation factor (*EF*1a) genes were selected for *I. accuminatus* (unpublished data). Six primer pairs representing different bacterial taxa and one eubacterial primer revealing the total bacterial population were used for qPCR assay (Supplementary Table 3). The specific bacterial primers were selected from previously published studies, while *Psedoxanthomonas* and *Serratia* genus-specific primers were designed in-house, based on 16S rRNA gene sequences available in NCBI. 10 µl of reaction mixture was prepared containing 4µl of gDNA (10 ng/µl), 5 µl SYBR® Green PCR Master Mix (Applied Biosystems), 0.5 µl forward and reverse primer (10 µM). Amplification conditions included initial denaturation at 95°C for 5 min, followed by 40 cycles of 95°C for 15 s and 60°C for 30 s. The relative quantification (RQ) of the selected bacterial population was estimated using the delta-delta Ct method (2^−ΔΔCt^), where ΔCt was estimated as the difference between the threshold Ct values with specific bacterial primers and the housekeeping reference gene. In this study, the 2^−ΔΔCt^ method reveals the fold change of the bacterial abundance relative to the housekeeping genes. It is worth mentioning here that the relative abundance of a specific bacterial population compared to the total bacterial population often lacks consistency (59). Hence, the housekeeping genes with stable gene expression were considered to normalize the data and determine the relative bacterial abundance. In the real-time analysis, the significant difference in relative bacterial abundance between different life stages of *I. sexdentatus* and *I. acuminatus* was estimated using the method described by Pekár & Brabec (60). A best linear model was elected under Akaike’s information criterion (AIC), and goodness of fit and heterogeneity were ensured by plotting residuals of the model against fitted values. Then, the variable was tested using ANOVA, and the difference between the categorical variable levels was compared by treatment contrasts t-test (61). All analyses were performed in the R 4.3.1 environment R (45).

## RESULTS

### Bacteriome structure

#### Sequencing statistics

The sequencing data from two pine bark beetle species, *I. sexdentatus* (ISX) and *I. acuminatus* (IAC), of different populations (lab-bred, wild collection) and life stages (larva, pupa, adult) along with host tissue (control wood, fed/gallery wood) generated 6,393,911 raw reads. A Phred Quality score >30 was used for quality control. Therefore, a total of 5,812,983 clean reads were obtained (Control wood-374,493; Fed wood-374,084; *I. sexdentatus* larvae-463,927; *I. sexdentatus* pupae-440,213; *I. sexdentatus* adult-446,527; *I. acuminatus* larvae-463,230; *I. acuminatus* pupae-446,017; *I. acuminatus* adult-372,076; *I. sexdentatus* wild adult-475,613; *I. sexdentatus* Control wood-364,782; *I. sexdentatus* Fed wood-381,812; *I. acuminatus* wild adult-447,180; *I. acuminatus* Control wood - 387, 312; *I. acuminatus* Fed wood-375,717) (Supplementary excel 1).

#### Bacterial relative abundance at distinct taxonomic levels

The bacterial sequences obtained from the two beetle species and wood samples generated a total of 15504 ASVs at a 97% similarity level (Supplementary excel 2). The Good’s coverage indicator (>99%) and the rarefaction curve indicated sampling comprehensiveness that represented the bacterial communities associated with the beetle and wood samples (Table 2, Supplementary figure 1). The predominant bacterial classes across all samples were Gammaproteobacteria, Alphaproteobacteria, Actinobacteria, Bacteroidia, Bacilli (Figure 1A). The relative abundance of Gammaproteobacteria was higher in *I. sexdentatus* pupae (92.8%) and adult (99.1%) samples compared to larvae (34.8%) and wild-collected adults (55.6%) samples (Supplementary excel 3). Similarly, the relative abundance of Gammaproteobacteria was highest in *I. acuminatus* larvae (95.1%) compared to other stages (82.9%, 79.4%) and wild-collected adults (85.2%). Furthermore, Alphaproteobacteria [*I. acuminatus* Control wood (11.3%)] and Gammaproteobacteria [*I. sexdentatus* Control wood (11.4%), *I. sexdentatus* Fed wood (69.5%), and *I. acuminatus* Fed wood (67%)] were the dominant class in different phloem wood samples (Supplementary excel 3).

**Figure 1:**
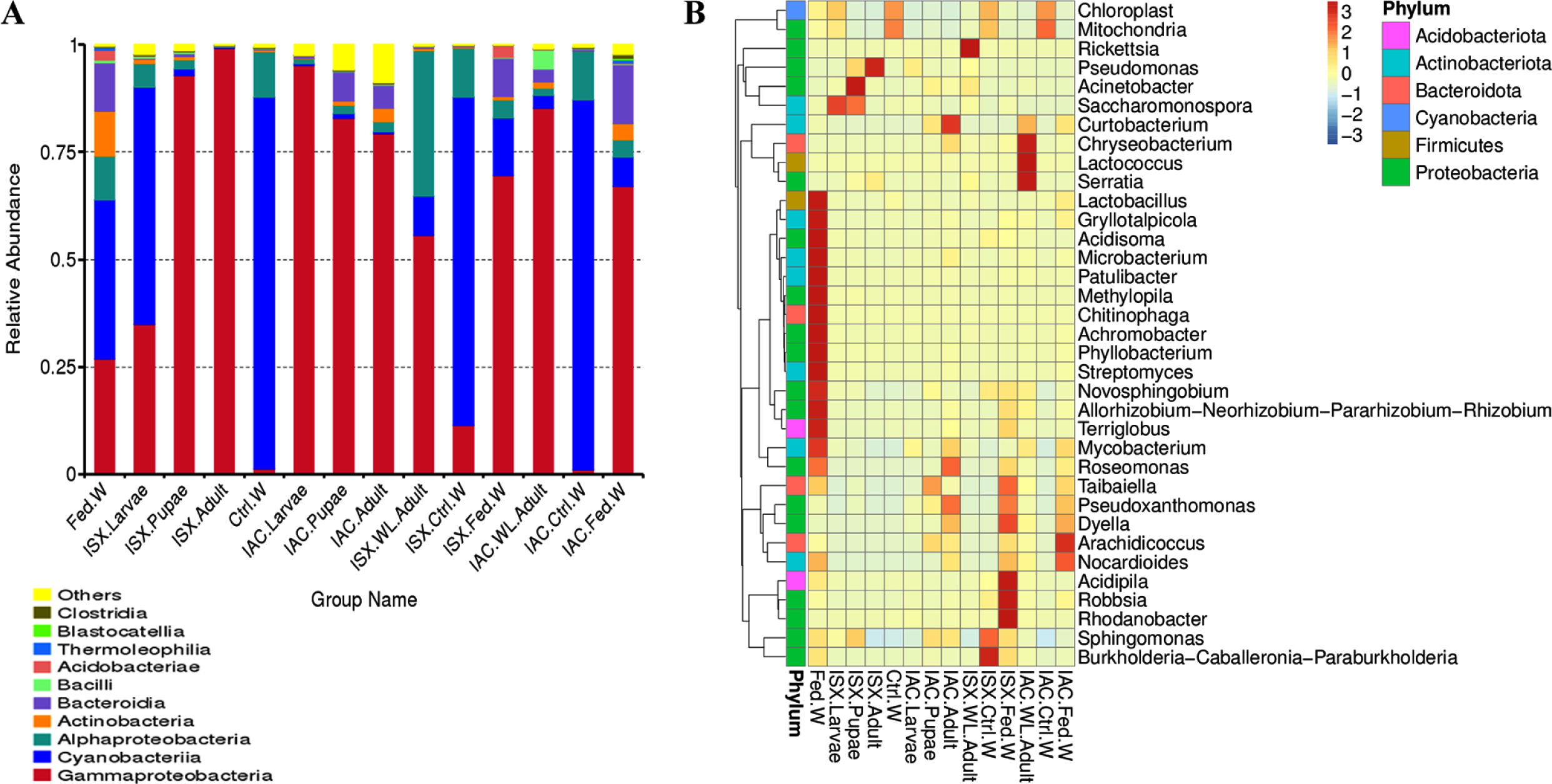
Bacterial diversity in lab-bred (life-stages), wild-type adult beetles and wood samples. (A) The bar plot represents the relative abundance of bacteriome at the class level (top 10). (B) Heatmap depicting the relative abundance of 35 dominant bacterial genera among lab-bred (life-stages), wild-type adult beetles and wood samples. The relative ASV abundance represented by a colour gradient where the darker colour indicates higher abundance, whereas the lighter colour indicates low abundance for a specific bacterial genus. (ISX-*Ips sexdentatus*, IAC-*Ips acuminatus*).

**Table 2:**
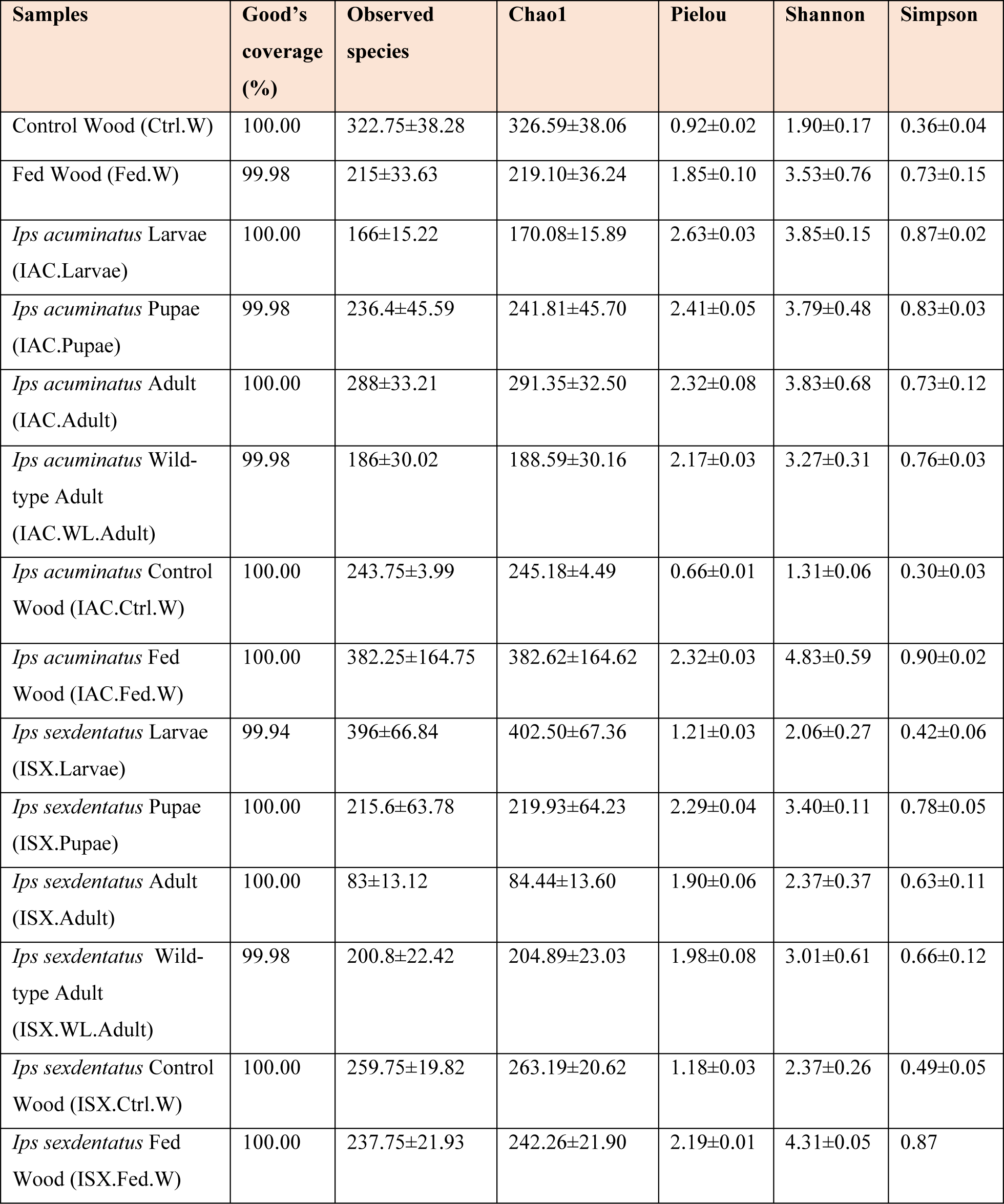
Alpha diversity indices. The data represented the mean value ±SE for five biological replicates across different life stages of two pine-feeding beetles and four biological replicates for the associated wood samples. (SE-Standard Error).

| Samples | Good's coverage (%) | Observed species | Chao1 | Pielou | Shannon | Simpson |
| --- | --- | --- | --- | --- | --- | --- |
| Control Wood (Ctrl.W) | 100.00 | 322.75±38.28 | 326.59±38.06 | 0.92±0.02 | 1.90±0.17 | 0.36±0.04 |
| Fed Wood (Fed.W) | 99.98 | 215±33.63 | 219.10±36.24 | 1.85±0.10 | 3.53±0.76 | 0.73±0.15 |
| *Ips acuminatus* Larvae (IAC.Larvae) | 100.00 | 166±15.22 | 170.08±15.89 | 2.63±0.03 | 3.85±0.15 | 0.87±0.02 |
| *Ips acuminatus* Pupae (IAC.Pupae) | 99.98 | 236.4±45.59 | 241.81±45.70 | 2.41±0.05 | 3.79±0.48 | 0.83±0.03 |
| *Ips acuminatus* Adult (IAC.Adult) | 100.00 | 288±33.21 | 291.35±32.50 | 2.32±0.08 | 3.83±0.68 | 0.73±0.12 |
| *Ips acuminatus* Wild-type Adult (IAC.WL.Adult) | 99.98 | 186±30.02 | 188.59±30.16 | 2.17±0.03 | 3.27±0.31 | 0.76±0.03 |
| *Ips acuminatus* Control Wood (IAC.Ctrl.W) | 100.00 | 243.75±3.99 | 245.18±4.49 | 0.66±0.01 | 1.31±0.06 | 0.30±0.03 |
| *Ips acuminatus* Fed Wood (IAC.Fed.W) | 100.00 | 382.25±164.75 | 382.62±164.62 | 2.32±0.03 | 4.83±0.59 | 0.90±0.02 |
| *Ips sexdentatus* Larvae (ISX.Larvae) | 99.94 | 396±66.84 | 402.50±67.36 | 1.21±0.03 | 2.06±0.27 | 0.42±0.06 |
| *Ips sexdentatus* Pupae (ISX.Pupae) | 100.00 | 215.6±63.78 | 219.93±64.23 | 2.29±0.04 | 3.40±0.11 | 0.78±0.05 |
| *Ips sexdentatus* Adult (ISX.Adult) | 100.00 | 83±13.12 | 84.44±13.60 | 1.90±0.06 | 2.37±0.37 | 0.63±0.11 |
| *Ips sexdentatus* Wild-type Adult (ISX.WL.Adult) | 99.98 | 200.8±22.42 | 204.89±23.03 | 1.98±0.08 | 3.01±0.61 | 0.66±0.12 |
| *Ips sexdentatus* Control Wood (ISX.Ctrl.W) | 100.00 | 259.75±19.82 | 263.19±20.62 | 1.18±0.03 | 2.37±0.26 | 0.49±0.05 |
| *Ips sexdentatus* Fed Wood (ISX.Fed.W) | 100.00 | 237.75±21.93 | 242.26±21.90 | 2.19±0.01 | 4.31±0.05 | 0.87 |

### Life stage-specific bacteriome

#### Bacterial associations in I. sexdentatus (ISX)

Our study revealed that the ASV distribution of the different life stages of both pine beetles comprises a pool of diverse bacterial populations. *I. sexdentatus* larvae, pupae, and adult stages contained 861, 355, and 126 unique ASVs, respectively (Figure 2A). Moreover, 61 ASVs were present in all life stages, constituting the core bacteriome. However, it is essential to mention that each ASV may not represent an individual species. The *I. sexdentatus* core consortium was categorized into 22 genera belonging to 22 families, where *Erwiniaceae*, *Pseudomonadaceae*, and *Yersiniaceae* were the dominating families, while *Pseudomonas and Serratia* were the prevalent bacterial genera (Supplementary excel 2, 4). The heatmap revealed the high abundance of *Pseudomonas* in adults, whereas *Acinetobacter, Saccharomonospora,* and *Rickettsia* were dominant in pupae, larvae, and wild adults, respectively (Figure 1B). The alpha diversity analysis showed that *I. sexdentatus* adult beetles had substantially lower bacterial richness (Chao1 84.44±13.60) than the other developmental stages (Chao1, larvae-402.50±67.36, p<0.01 and pupae-219.93±64.23, p<0.05) (Table 2, Supplementary figure 2, Supplementary excel 5). However, *I. sexdentatus* pupae represented significantly higher bacterial diversity (Shannon-3.40±0.11 and Simpson-0.78±0.05) compared to the larvae (Shannon-2.06±0.27, p<0.01 and Simpson-0.42±0.06, p<0.05) (Table 2, Supplementary figure 2). Similarly, the bacterial community evenness was significantly higher in *I. sexdentatus* pupae (2.29±0.04) compared to larvae (1.21±0.03) (p<0.01) (Table 2, Supplementary figure 2, Supplementary excel 5). Subsequently, metastat analysis revealed the differential abundance of bacterial genera (top 10) between different developmental stages (Table 3). Among the bacterial genera present, *Pseudomonas* was the most dominant bacterial genus in all the three life stages of *I. sexdentatus* beetles, with differences in their relative abundance. LEfSe represented the key bacterial biomarkers in *I. sexdentatus* life stage-specific bacterial populations (Figure 2D, Supplementary figure 3A). *I. sexdentatus* pupae documented the bacterial families, including *Enterobacteriaceae*, *Erwiniaceae*, and *Moraxellaceae,* as biomarkers (Figure 2D). In contrast, *Yersiniaceae* and *Pseudomonadaceae* were the biomarkers of *I. sexdentatus* adults. Similarly, *I. sexdentatus* larvae represented the class Alphaproteobacteria as a distinct biomarker.

**Figure 2:**
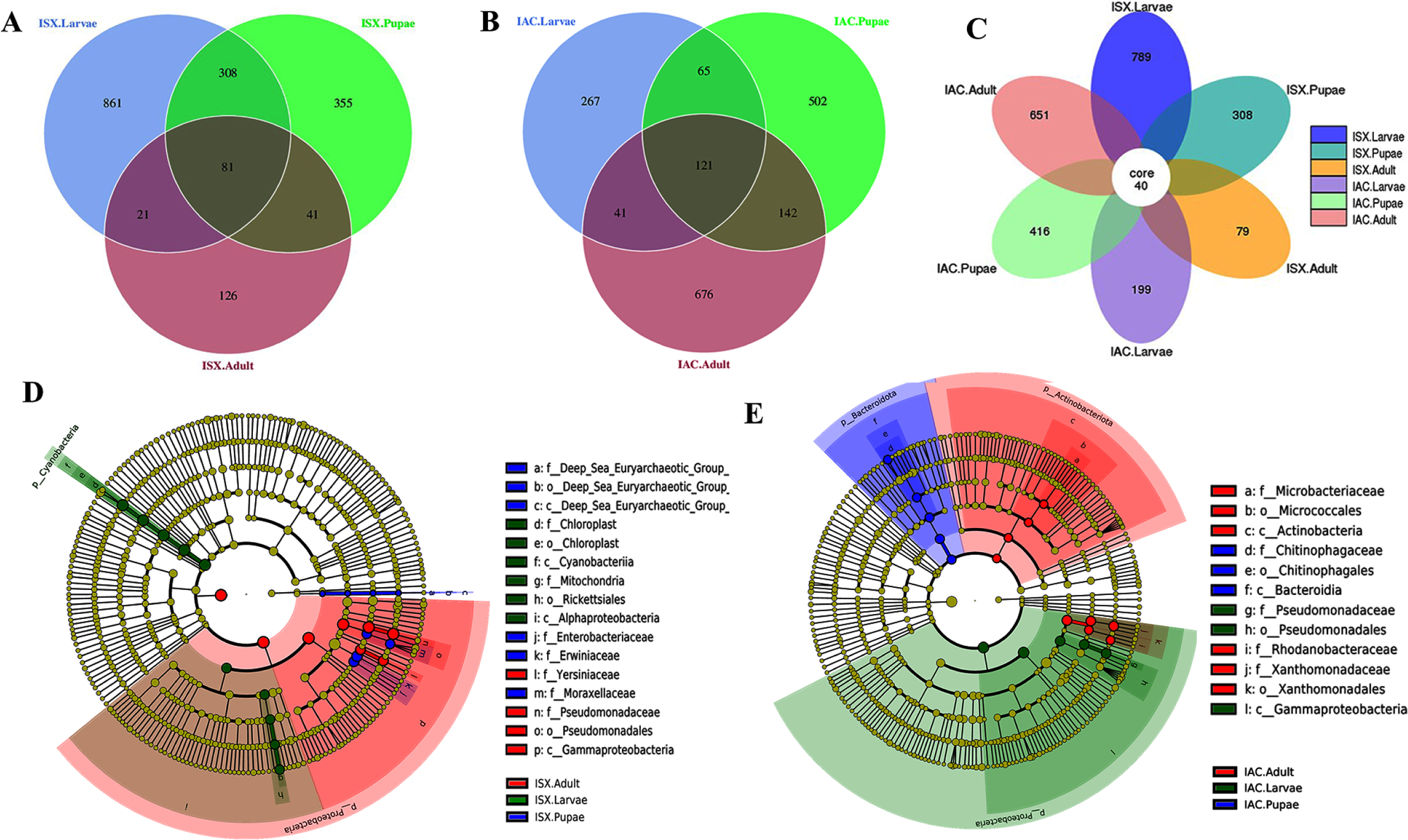
Core bacteriome. (A) Venn diagram illustrating bacterial ASV distribution in *I. sexdentatus* life stages (ISX.Larvae, ISX.Pupae, ISX.Adult). (B) Venn diagram depicting bacterial ASV distribution in *I. acuminatus* life stages (IAC.Larvae, IAC.Pupae, IAC.Adult). (C) Flower diagram representing the core ASVs across the life stages of two pine beetles. (D) Cladogram illustrating the results from LefSe analysis revealing the biologically consistent, statistically significant bacterial biomarkers across different life stages of *I. sexdentatus* (ISX). (E) Cladogram representing significantly distinct bacterial biomarkers across *I. acuminatus* (IAC) life stages. Distinct taxonomic level (phylum to genus) is denoted in the circle from inward to outward. The different coloured nodes (red, green and blue) represent bacterial species that play a significant role in different life stages across two different beetles (ISX and IAC), whereas yellowish-green circles represent non-significant bacterial species. Specific bacterial biomarkers are denoted by letters above the circles. The size of the nodes represents the relative abundance of the bacterial species at a particular taxon. (ISX-*Ips sexdentatus*, IAC-*Ips acuminatus*).

**Table 3:**
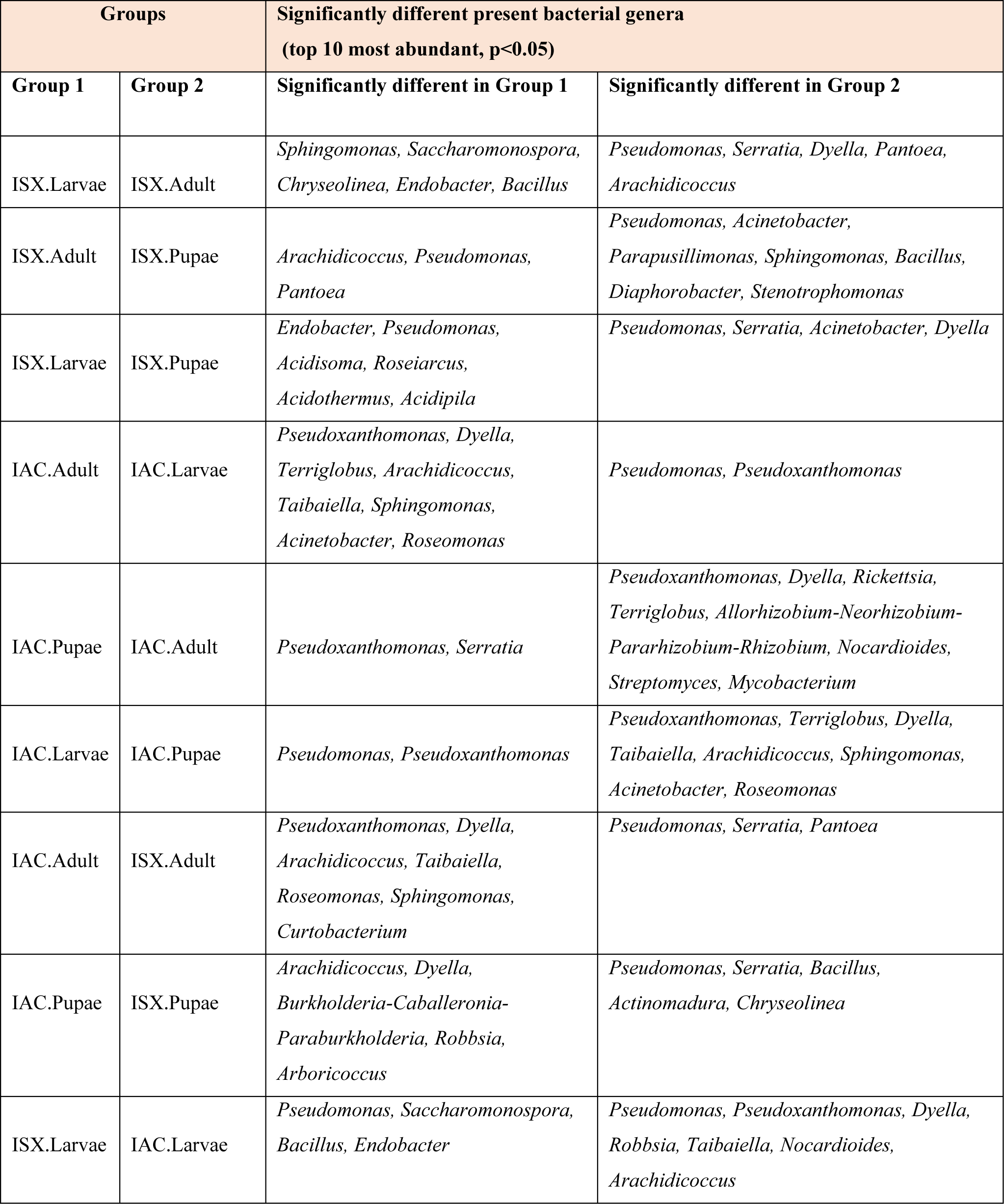
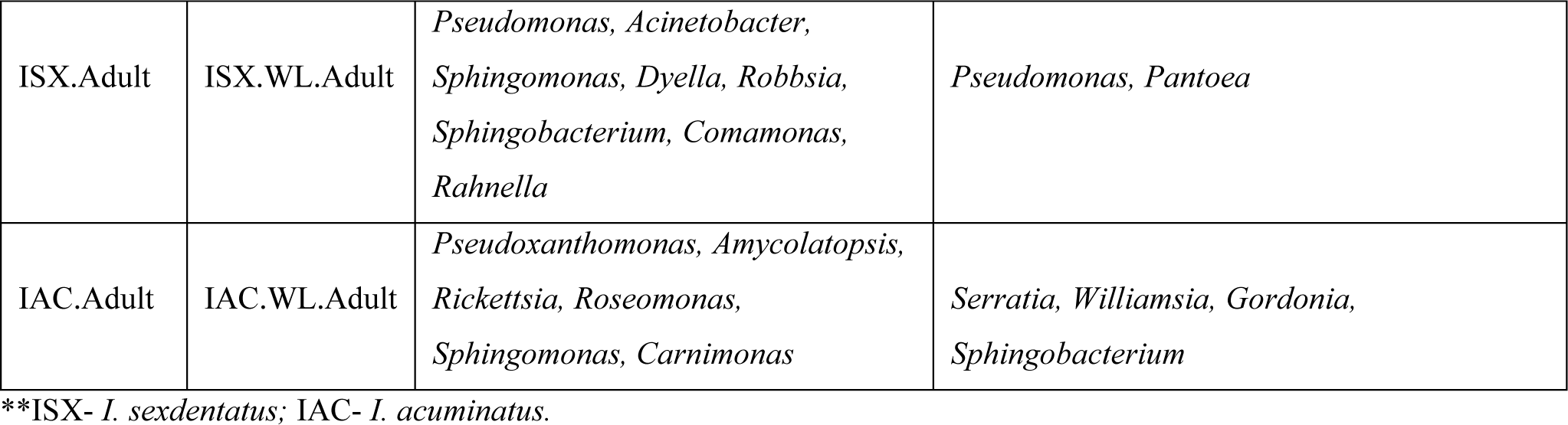
Metastat analysis representing the top 10 differently abundant bacterial genera across different life stages of *Ips* pine beetles.

#### Bacterial associations in I. acuminatus (IAC)

The core bacteriome in *I. acuminatus* comprised 121 ASVs that were categorized into 28 families and 35 genera (Figure 2B), with *Erwiniaceae*, *Pseudomonadaceae,* and *Rhodanobacteraceae* being the most dominant bacterial families (Supplementary excel 2, 4). Comparing between the developmental stages, *Mycobacterium* and *Pseudomonas* showed a high abundance in *I. acuminatus* larvae, while *Taibaiella, Sphingomonas, Arachidicoccus,* and *Curtobacterium* were dominant in the pupal stage (Figure 1B). Furthermore, the alpha diversity indices revealed higher bacterial richness in *I. acuminatus* adults (Chao1, 291.35±32.50) compared to larvae (Chao1, 166±15.22) (p<0.05) (Table 2, Supplementary figure 2, Supplementary excel 5). However, no stage-specific differences in bacterial diversity and evenness were observed in *I. acuminatus* beetles (Table 2). Metastat analysis revealed a significantly high abundance of *Pseudomonas* in *I. accuminatus* larvae, while *Pseudoxanthomonas* was prevalent in adults and pupae (Table 3). Furthermore, LEfSe analysis corresponds to the bacterial biomarkers in *I. acuminatus* life stages (Figure 2E, Supplementary figure 3B). *I. acuminatus* pupal biomarkers were categorized into phylum Bacteroidota and class bacteroidia (Figure 2E), while members from the phyla Proteobacteria and Gammaproteobacteria were represented as the biomarkers in *I. acuminatus* larvae. Similarly, *I. acuminatus* adults documented Proteobacteria (family-*Rhodanobactericeae*, *Xanthomonadeceae*) and Actinobacteria (family-*Microbactericeae*) as predominant biomarkers.

#### Comparing bacteriome of two pine-feeding Ips beetles

Comparing the two beetle species, *I. acuminatus* and *I. sexdentatus,* 40 ASVs were shared across the developmental stages in both species (Figure 2C) that were assigned to 15 families encompassing different genera like *Pseudoxanthomonas, Pseudomonas, Serratia, Dyella, Taibaiella, Acinetobacter, Sphingomonas, Allorhizobium-Neorhizobium-Pararhizobium-Rhizobium, Acidipila, Arboricoccus, Saccharomonospora, Roseomonas, Acidisoma, Cutibacterium, Providencia* (Supplementary excel 6). Furthermore, the adult and larval stages of both species (*I. sexdentatus* and *I. acuminatus*) showed a significant difference (p<0.01) in bacterial richness. Although there was not much variation in bacterial diversity and evenness within the pupal and the adult stages between the two beetle species (*I. acuminatus* and *I. sexdentatus*), *I. acuminatus* larvae showed significantly higher bacterial diversity (Shannon-3.85±0.15, Simpson-0.87±0.02) compared to *I. sexdentatus* larvae (Shannon-2.06±0.27, Simpson-0.42±0.06), p<0.01) (Table 2, Supplementary figure 2, Supplementary excel 5). Subsequently, *I. acuminatus* larvae (2.63±0.03) represented more significant bacterial evenness than *I. sexdentatus* larvae (p<0.01). Additionally, NMDS using unweighted UniFrac distances revealed the differences between bacterial communities of *I. sexdentatus* and *I. acuminatus* by hierarchically clustering different life stages and wood samples (Figure 3A). The larval and adult beetle-associated bacteria in *I. acuminatus* and *I. sexdentatus* had significant differences within and among them.

**Figure 3:**
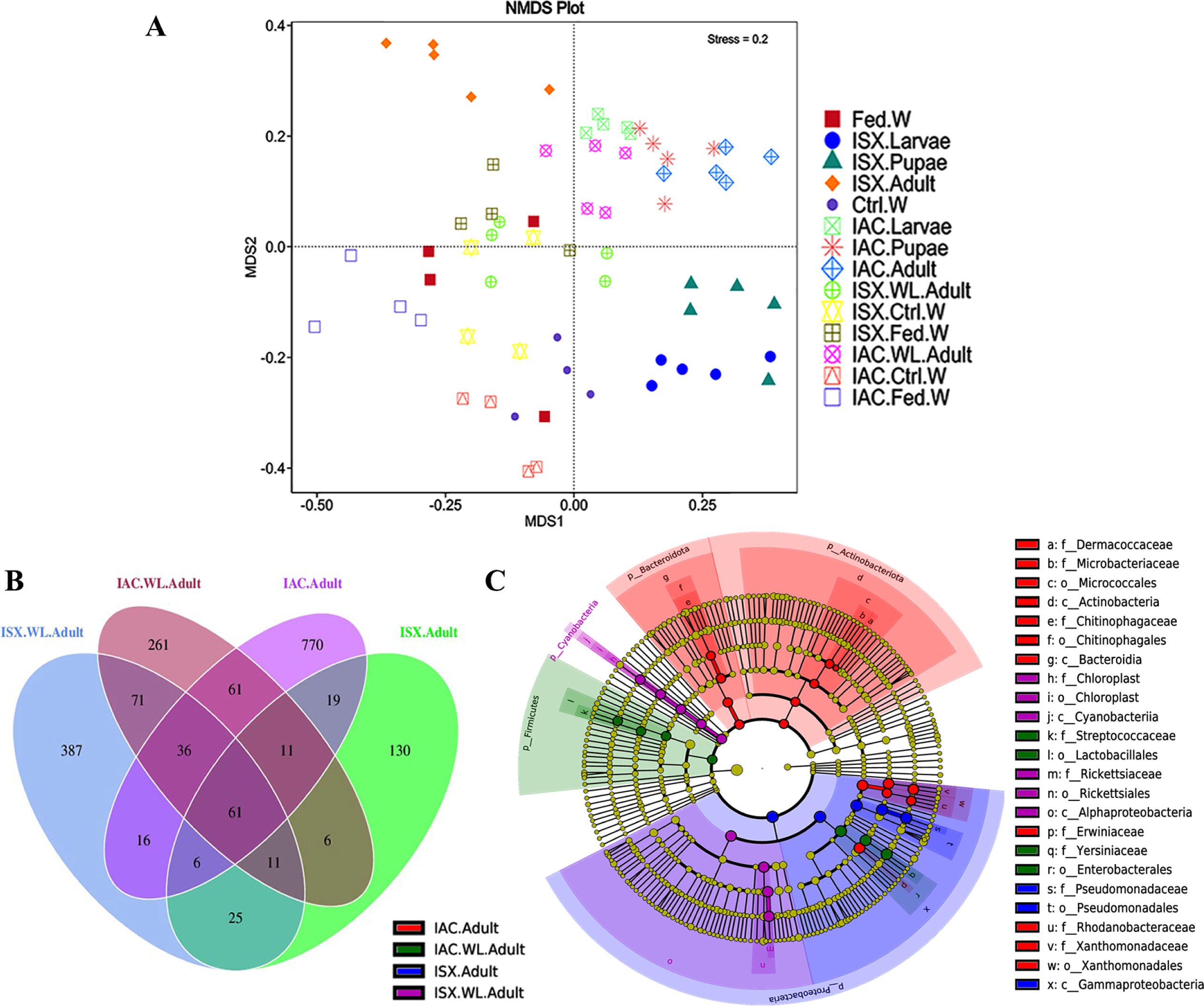
Impact of lab breeding on bacterial association. (A) Flower diagram representing the shared and unique ASVs in *I. sexdentatus, I. acuminatus* wild collected adults (ISX.WL.Adult, IAC.WL.Adult) and lab-bred adult beetles (ISX.Adult, IAC.Adult). (B) Cladogram representing significant bacterial biomarkers among *I. sexdentatus, I. acuminatus* wild type and lab-bred beetles. (ISX-*Ips sexdentatus*, IAC-*Ips acuminatus*).

### Impact of beetle lab breeding on bacterial assemblage

#### I.sexdentatus: wild vs lab-bred

According to our study, the lab-bred adult population (F2 generation) and the wild beetle population have considerable differences in the composition of their ASVs. The *I. sexdentatus* wild and lab-bred adults had 387, 130 unique ASVs, and 103 shared ASVs (Figure 3B). The unique ASVs in wild adults accounted for 76 families and 82 genera, including *Erwiniaceae* and *Pseudomonadaceae* as the dominant families (Supplementary excel 7), while *Serratia* and *Pseudomonas* were the highly abundant genera. Similarly, in lab-bred adults (F2, lab-breed), bacterial ASVs were categorized into 40 genera belonging to 40 families, having *Erwiniaceae* and *Pseudomonadaceae* as dominant families, whereas *Pantoea and Pseudomonas* were the prevalent genera (Supplementary excel 7). Alpha diversity comparisons revealed that in *Ips sexdentatus* samples, the adult beetles (84.44±13.60) had lower bacterial richness than wild adults (204.89±23.03, p<0.01) (Table 2, Supplementary figure 2, Supplementary excel 5). Moreover, metastat analysis revealed significant differences in bacterial abundance between the lab-bred and wild-collected beetles (Table 3). Furthermore, the lab-bred and wild-collected beetles possess distinct bacterial markers (Figure 3C, Supplementary figure 3C). For instance, LEfSe analysis revealed the members of the class Gammaproteobacteria (family-*Pseudomonadaceae*) as the biomarkers of lab-bred adults, while Alphaprotobacteria (family-*Rickettsiaceae*) was represented as the biomarker of wild-collected adults (Figure 3B).

#### I. acuminatus: wild vs lab-bred

The lab-bred and wild-collected *I. acuminatus* adult beetles comprised a higher number of unique bacterial ASVs (770 and 261) than shared ones (169) (Figure 3B). Lab-bred adult bacteriome included 53 families and 73 genera (Supplementary excel 7), where *Acidobacteriaceae*_(Subgroup_1), *Corynebacteriaceae*, *Microbacteriaceae* were highly abundant bacterial families (Supplementary excel 7). In comparison, *I. acuminatus* wild-collected adult population comprised 45 families and 58 genera, including *Xanthomonadaceae*, *Erwiniaceae*, and *Microbacteriaceae* as dominating families (Supplementary excel 7). The heatmap indicated *Curtobacterium, Roseomonas,* and *Pseudoxanthomonas* dominated lab-bred adults, while *Cryseobacterium, Lactococcus,* and *Serratia* were prevalent in wild *I. acuminatus* beetles (Figure 1B). Furthermore, the alpha diversity analysis revealed that lab-bred *I. acuminatus* adult (Chao1 291.35±32.50) had higher bacterial richness compared to wild adults (188.59±30.16) (p<0.05) (Table 2, Supplementary figure 2). However, no significant differences in bacterial diversity were observed (Table 2, Supplementary figure 2, Supplementary excel 5). LEfSe analysis represented biomarkers in *I. accuminatus* adult belonging to different phyla-Actinobacterioda (family-*Dermacoccaceae*, *Microbacteriaceae*; order-Micrococcales; class-Actinobacteria), Bacteroidota (family-*Chitinophagaceae*, order-chitinophagales, class-bacteroidia), Proteobacteria (family-*Rhodanobactericeae*, *Xanthomonadeceae*, *Erwiniaceae*) (Figure 3C, Supplementary figure 3C). *I. acuminatus* wild adults contained biomarkers belonging to two phyla-Proteobacteria (family-*Yersiniaceae*; order-Enterobacterales) and Firmicutes (family-*Streptococcaceae*; order-Lactobacillales). Nevertheless, the wild adult beetles for both *Ips* species were clustered separately from lab-bred adults in the NMDS plot, indicating laboratory breeding impact on beetle bacteriome (Figure 3A).

### Host contribution in shaping beetle bacteriome

Comparing the *I. sexdentatus* wild beetles with their control and fed woods revealed a consortium of 121 shared ASVs (Figure 4A). The *I. sexdentatus* gallery wood (32 bacterial families and 28 bacterial genera) shared more ASVs than the unfed control (17 bacterial families, 13 bacterial genera) (Supplementary excel 8), which is probably due to beetle feeding. The shared bacteriome documented a high abundance of bacterial families *Pseudomonadaceae, Yersiniaceae, and Erwiniaceae,* while *Pseudomonas* is the dominant genus. The unfed pine wood (*I. sexdentatus* Control wood) from the forest documented a high abundance of bacterial genera-*Burkholderia _Caballeronia_Paraburkholderia* and *Sphingomonas,* whereas *Acidipila, Dyella, Robbsia,* and *Rhodanobacter* were dominant in gallery wood of the wild *I. sexdentatus* (Figure 1B).

**Figure 4:**
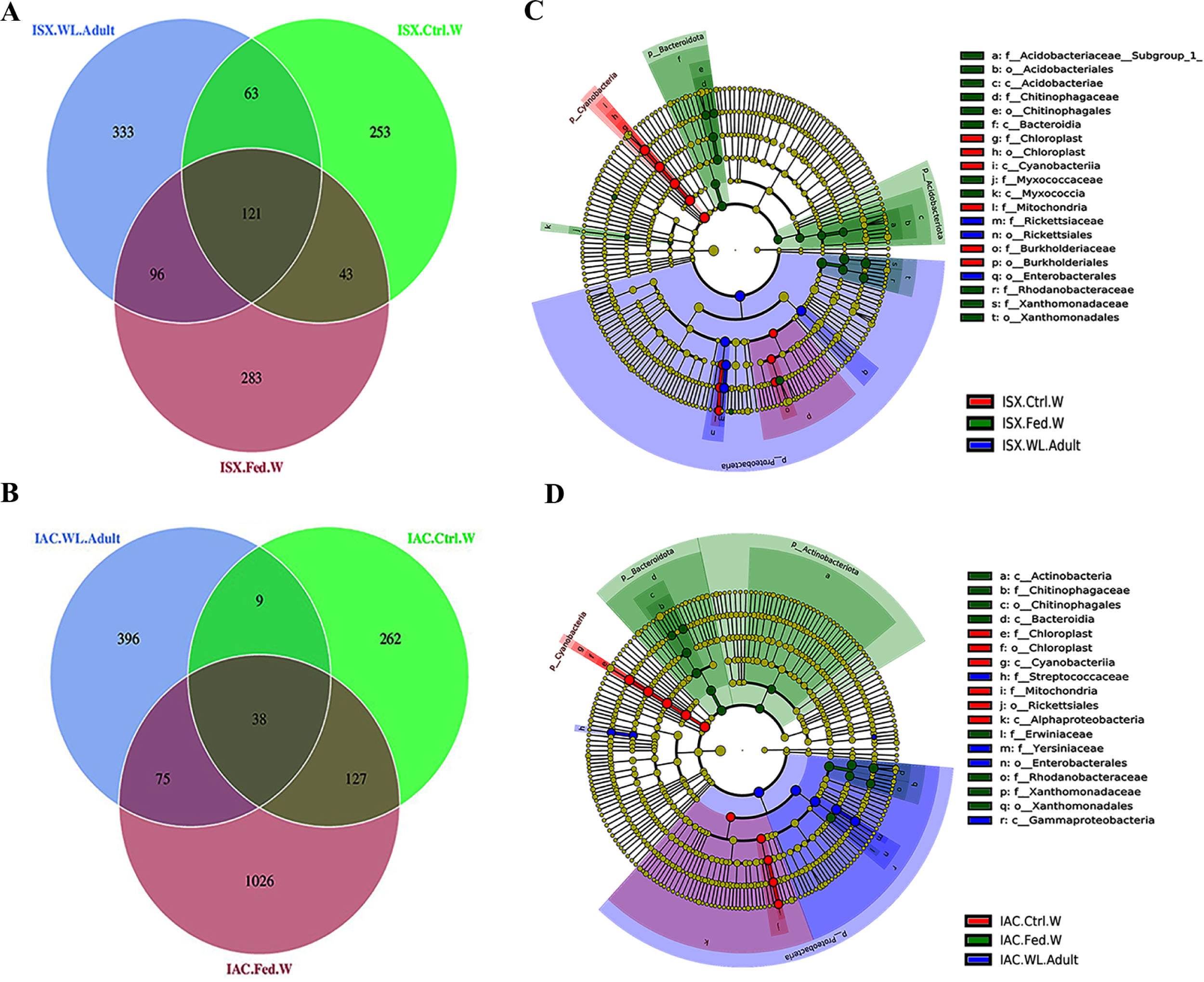
Host contribution in shaping beetle bacteriome. (A) Venn diagram showing the bacterial ASVs contribution of wood bacteriome in shaping *I. sexdentatus* bacteriome. (B). Venn diagram showing the bacterial ASVs contribution of wood bacteriome in shaping *I. acuminatus* bacteriome. (C) Non-metric Multi-Dimensional Scaling (NMDS) based on unweighted UniFrac distance matrix represents bacterial diversity variation in different life stages of two beetles along with the wood samples. (D) Cladogram representing significant bacterial biomarkers in *I. sexdentatus* wild-adult beetle and different wood types (control wood and fed wood). (E) Cladogram representing significant bacterial biomarkers among *I. acuminatus* wild adult beetle and different wood types (control wood and fed wood). (ISX-*Ips sexdentatus*, IAC-*Ips acuminatus*).

Similarly, the *I. acuminatus* wild beetles and their fed wood revealed a higher proportion of shared ASV (30 families, 39 genera) population than the control wood (6 families and 6 genera) (Figure 4B, Supplementary excel 8). Subsequently, the shared bacterial community between *I. acuminatus* Control wood and *I. acuminatus* fed wood was dominated by 4 families (*Erwiniaceae*, *Yersiniaceae*, *Weeksellaceae*, *Streptococcaceae*) and 3 genera (*Serratia, Chryseobacterium, Lactococcus*). *Pseudoxanthomonas, Arachidicoccus, Dyella,* and *Nocardioides* were prevalent in gallery wood of the wild *I. acuminatus*, which in turn had a lower abundance in unfed control pine wood (*I. acuminatus* Control wood), indicating an alteration in pine wood bacterial assemblage after beetle feeding (Figure 1B).

Several biomarkers were found while comparing the adult and wood bacterial communities. Proteobacteria was the significant biomarker in the control wood (*I. sexdentatus* and *I. acuminatus* control wood) and adult (*I. sexdentatus* and *I. acuminatus* wild adult) population (Figure 4C and 4D, Supplementary figure 3D and 3E). However, at other taxonomic levels, the biomarkers differed. *I. sexdentatus* control wood biomarkers belong to the family-*Burkholderiaceae*, order-Burkholderiales, whereas *I. accuminatus* control wood biomarkers belong to order-Rickettsiales, class-Alphaproteobacteria. Subsequently, *I. sexdentatus* wild adults’ biomarkers were categorized into the family-*Rickettsiaceae*, order-Rickettsiales, Enterobacterales, and the biomarkers for *I. acuminatus* wild adults were classified into the family-*Streptococcaceae*, *Yersiniaceae*, order-Enterobacterales, class-Gammaproteobacteria. However, in the fed wood samples from two beetle species (*I. sexdentatus* and *I. acuminatus* fed wood), Proteobacteria, Acidobacteriota, and Bacterodiota were the predominant phyla. Similarly, there were differences at other taxa levels. *I.sexdentatus* fed-wood biomarkers belonged to families such as *Acidobacteriaceae* _(Subgroup_1), *Chitinophagaceae*, *Myxococcaceae*, *Rhodanobacteraceae*, and *Xanthomonadaceae* (Figure 4C). Similarly, *Chitinophagaceae*, *Erwiniaceae*, *Rhodanobacteraceae*, and *Xanthomonadaceae* were the prevalent biomarkers in *I. acuminatus* fed wood (Figure 4D). Investigating the functional relevance of these biomarkers in different sample groups will be intriguing.

We observed considerable differences in bacterial diversity and community evenness in different wood samples. However, no significant difference in bacterial richness was observed between wood samples (Table 2, Supplementary figure 2, Supplementary excel 5). The fed/gallery wood samples collected from the forest demonstrated significantly different higher bacterial community diversity (*I. sexdentatus* Fed wood, Shannon-4.31±0.05, Simpson-0.87; *I. acuminatus* Fed wood, Shannon-4.83±0.59, Simpson-0.90±0.02) and evenness (*I. sexdentatus* Fed wood-2.19±0.01; *I. acuminatus* Fed wood-2.32±0.03) than their respective control wood samples (*I. sexdentatus* Control wood Shannon-2.37±0.26, Simpson-0.49±0.05, evenness-1.18±0.03; *I. acuminatus* Control wood Shannon-1.31±0.06 and Simpson-0.30±0.03, evenness-0.66±0.01) (p<0.05) (Table 2, Supplementary figure 2, Supplementary excel 5). Such findings indicate the enrichment of bacterial species in the feeding gallery after beetle feeding.

Consequently, the NMDS plot represented *I. acuminatus* wild adults, fed wood, and control wood into distinct clusters (Figure 3A). However, no such clustering was observed in the case of wild-collected *I. sexdentatus* samples. In contrast, comparing lab-bred and wild-collected wood samples (control wood and fed wood) for the *I. sexdentatus* revealed a distinct bacterial population, suggesting the influence of environment and beetle feeding as drivers in shaping the host microbiome (Figure 3A, Supplementary figure 4, Supplementary excel 9).

### Relative bacterial quantification using quantitive PCR (qPCR) assay

#### Comparing *I. sexdentatus* life stages

The qPCR assay revealed a relatively high abundance of total bacterial population (eubacterial primers) in *I. sexdentatus* adults compared to all other life stages (Supplementary figure 5A, Supplementary excel 10). Precisely, there is a difference between the life stages of *I. sexdentatus* (ANOVA; df=2, p < 0.01). However, the relative bacterial abundance between larval and pupal stages is marginally varied (contrast t-test; p = 0.099); adult bacterial assemblage was significantly different from larvae (contrast t-test; p < 0.01) but slightly diverse from pupae (contrast t-test; p= 0.054). The relative abundance of Bacteroidetes (phylum), *Enterobacteriaceae*(Family)*, Pseudoxanthomonas, Pseudomonas, and Serratia* (genus) varied within the life stages of *I. sexdentatus* (Figure 5A,5B,5D,5E,5F). No abundance difference was observed for Firmicutes (phylum) (Figure 5C).

**Figure 5:**
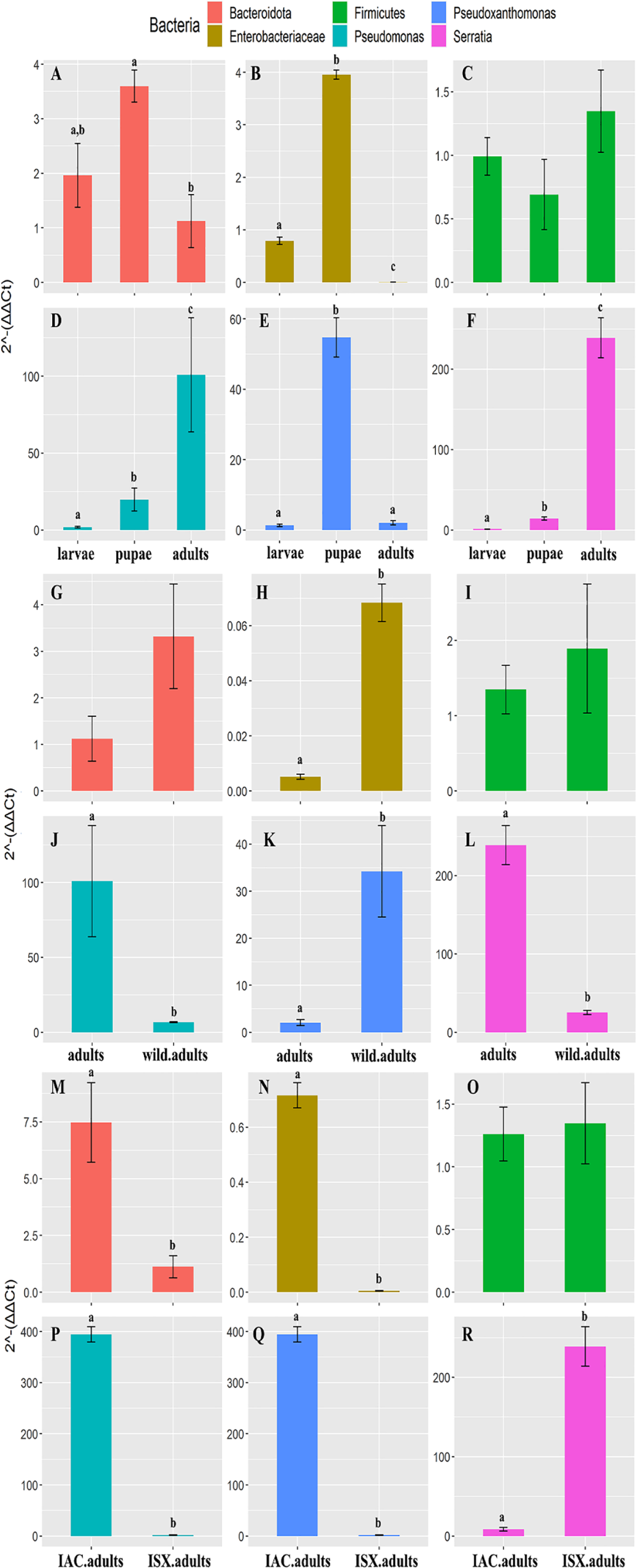
Quantitative PCR assay representing the relative abundance of selected bacterial taxa present in the two pine-feeding beetles. The 2^−ΔΔCt^ revealed the fold change of the bacterial abundance relative to the stable reference genes. The variable was tested using ANOVA, and the difference between the categorical variable levels was compared by treatment contrasts t-test. The level of statistical significance was represented by letters. A-F) *I. sexdentatus* life stage comparisons; G-I) *I. sexdentatus* wild vs. bred adults; M-R) *I. sexdentatus vs I. accuminatus* adults.

#### Wild ve lab-bred

Similarly, the overall difference in bacterial abundance (eubacterial primers) between wild and breeding adult beetles was non-significant (ANOVA; df=1, p = 0,076) (Supplementary figure 5B, Supplementary excel 10). However, the lab-bred *I. sexdentatus* adults showed a higher abundance of *Pseudomonas* and *Serratia* than wild adults (Figure 5J, 5L). Whereas *Enterobacteriaceae*, *Pseudoxanthomonas* has a higher abundance in wild beetles (Figure 5H,5K). No abundance differences were observed for Bacteroidetes and Firmicutes (Figure 5G, 5I).

#### I. sexdentatus vs I. accuminatus

Comparing the two pine beetles, adults of *I. acuminatus* had significantly higher relative abundance than *I. sexdentatus* (ANOVA; df=1, p < 0.001)(Supplementary figure 5C, Supplementary excel 10). The relative abundance of Bacteroidetes (phylum), *Enterobacteriaceae* (Family), *Pseudomonas*, and *Pseudoxanthomonas* (genus) was significantly higher in *I. acuminatus* adults (Figure 5M, 5N,5P,5Q). However, *Serratia* (genus) abundance was greater in *I. sexdentatus* adults (Figure 5R). No significant differences in the relative abundance of Firmicutes were observed for any comparisons (Figure 5C,5I,5O).

### Functional prediction of the pine beetle bacteriome

The PICRUSt2 predicted the probable functional profile of the bacterial communities in the two pine-feeding beetles based on the relative abundance of 16S rRNA gene sequences. Individual bacterial ASVs were attributed to a gene function predicting the functional diversity based on different databases (KEGG and COG) and represented in the heatmap cluster (Supplementary Figure 6). For instance, the functional prediction based on KEGG database represented a high abundance of MFS transporter, type IV secretion system protein VirB6, type I restriction enzyme M protein, mRNA interferase RelE/StbE in *I. sexdentatus* wild adult compared to other life stages (Supplementary figure 6A). Similarly, the iron complex transport system substrate-binding protein, methyl-accepting chemotaxis protein, and polar amino acid transport system permease protein had relatively higher abundance in *I. acuminatus* larvae and pupae than in adults (Supplementary figure 6A). ASV comparison with the COG database revealed a higher abundance of Flavin-dependent oxidoreductase in *I. acuminatus* larvae and pupae (Supplementary figure 6B). Alternatively, glutathione S-transferase, DNA-binding protein (AcrR family), and DNA-binding transcriptional regulator (LysR family) had comparatively high abundance in the adults (Supplementary figure 6B). Interestingly, N-acetyltransferase and glutathione S-transferase (GST) are highly present in the pupal stage of both beetles (Supplementary figure 6B). Such observations suggest that genes showing relatively similar abundance might execute fundamental functions in the beetle holobiont and constitute the core bacteriome, while a stage-specific abundance of specific genes can indicate stage-specific function.

We also illustrated the functional profile based on the significant bacterial abundance and genes coding for enzymes (represented by EC number) in two pine-feeding beetles and generated comparative heatmaps based on three fundamental functions such as amino acid biosynthesis, cell wall degradation, vitamins, and co-factors biosynthesis (Figure 6, 7, Supplementary figure 7, 8, Supplementary excel 11,12). Genes involved in amino acid biosynthesis, such as *leu C* and *leu D,* were highly abundant in *I. sexdentatus* larvae, while the abundance of gene *dap A* was higher in pupae and adults (Figure 6) (abbreviation table contains full names). While in *I. acuminatus*, the genes *leu C* and *leu D* were highly present in both larvae and pupae compared to the adults. Subsequently, *argD, and dapC* were abundant in the larvae, whereas *metH*, *trp GD, trpA* were highly present in *I. acuminatus* lab-bred adults (Figure 7). Furthermore, genes like *ALDO*, *rpe* involved in cell wall biosynthesis documented higher abundance in *I. sexdentatus* larvae and pupae, respectively (Figure 6). On the contrary, *chiA* and *pcaC* had higher abundance in *I. sexdentatus* wild and lab-bred adult samples (Figure 6). In *I. acuminatus, pcaC*, and *HEXA_B* were documented in higher abundance in larval and pupal stages, respectively, whereas nagA and *xylB* were significantly abundant in the lab-bred adult samples (Figure 7). Similarly, *I. sexdentatus* pupae and adult stage showed a higher abundance of *fabG,* contributing to the biosynthesis of vitamins and co-factors (Figure 6).

**Figure 6:**
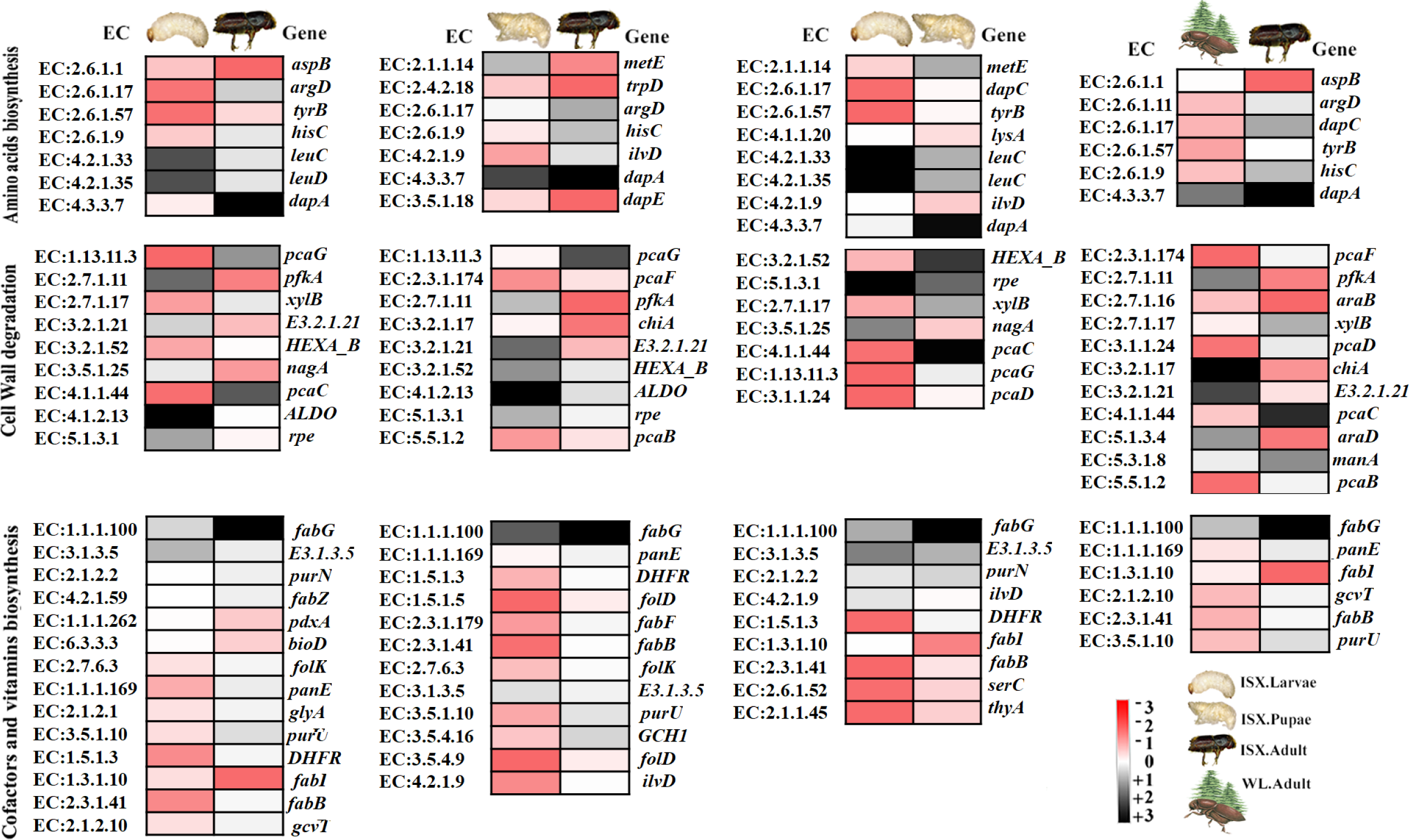
PICRUSt2 results for *I. sexdentatus*. Summarized heatmaps representing highly different and significantly abundant (p<0.05, t-test) bacterial genes coding for enzymes involved in amino acid biosynthesis, cell wall degradation, vitamins and co-factors biosynthesis among various life stages of ISX beetles and wild-type ISX. (ISX-*Ips sexdentatus*).

**Figure 7:**
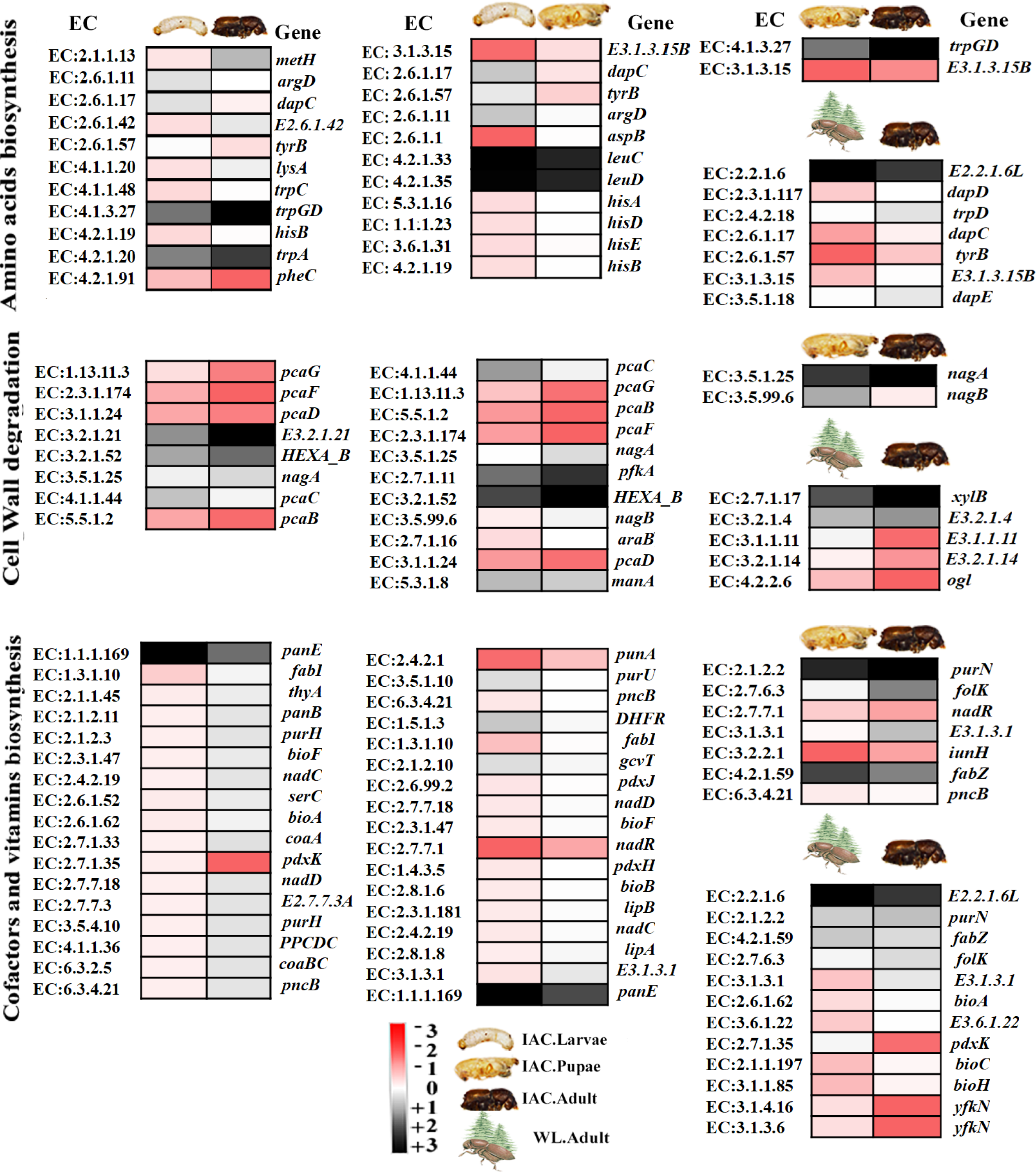
PICRUSt2 results for *I. accuminatus*. Summarized heatmaps representing highly different and significantly abundant (p<0.05, t-test) bacterial genes coding for enzymes involved in amino acid biosynthesis, cell wall degradation, vitamins and co-factors biosynthesis among various life stages of IAC beetles and wild-type IAC. Darker colours represent higher abundance, whereas lighter colours indicate a lower abundance of a specific gene/enzyme. (IAC-*Ips acuminatus*).

Finally, to visualize the bacterial abundance in relation to the PICRUST2 data, a summary figure was illustrated representing the top 5 bacterial families and categorizing significantly abundant enzymes present in different stages of both pine-feeding beetles (Figure 8). The categorization is based on the 4-fold change in enzyme abundance between two stages at different levels of significance (p<0.001 designated as “***”, p<0.01 denoted by “**”, p<0.05 designated as “*”) (Supplementary excel 13-15).

**Figure 8:**
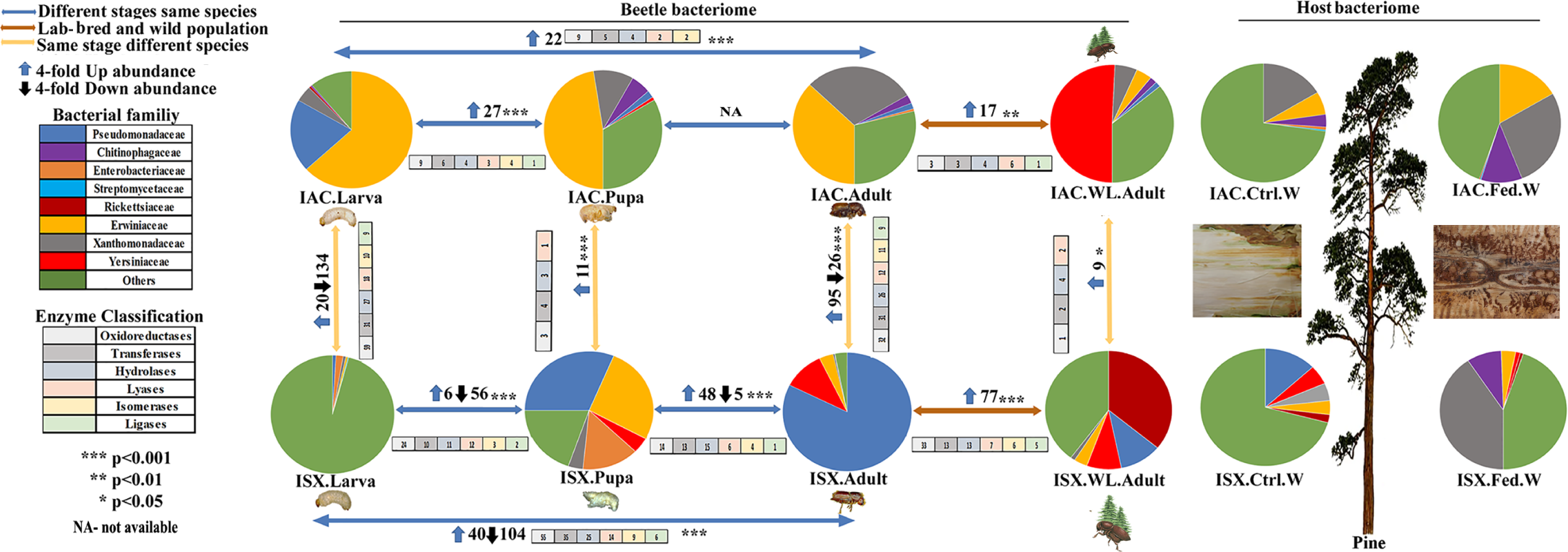
Schematic diagram containing the top 5 bacterial families at each stage of pine beetles and their respective wood samples (control wood, gallery wood). Different coloured arrows represent a different comparison level (within a beetle species, between two species, lab-bred and wild species, between wild species). The up and down arrows represent a 4-fold change in potential bacterial EC abundance at a different level of significance (p<0.001 designated as “***”, p<0.01 designated as “**”, p<0.05 designated as “*”). The coloured boxes with numbers indicate different enzymatic classes present in each comparison. (ISX-*Ips sexdentatus*, IAC-*Ips acuminatus*).

## DISCUSSION

Conifers have evolved a formidable defence against pests and pathogens (16, 62). In contrast, pests, including bark beetles and their associated microbiota, can successfully invade host trees by compromising host defence (25). The insect microbiota is often influenced by the diet, sex, life stages, and the environment and facilitates the expansion of the ecological and evolutionary potential of their hosts (63, 64). There has been growing interest in understanding the role of insect-microbial association in shaping the plasticity of insects in recent times. For instance, a recent study on Eurasian spruce bark beetles, *I. typographus*, reported the developmental stage and the geo-location as drivers in shaping the beetle-microbial association (27–29). Nevertheless, the gut bacterial dynamics in the pine beetle *Dendroctonus rhizophagus* revealed that the presence of persistent bacterial communities across the life stages of the beetle might be essential for ensuring certain physiological functions for the host (65). However, comprehensive information on the contribution of life stage and environment on the pine-feeding *Ips* bark beetles, *I. sexdentatus,* and *I. acuminatus* (Coleoptera: Curculionidae) is lacking. Hence, the present study is focused on the influence of life stage and environment on the bacterial communities associated with the two pine-feeding *Ips* bark beetles.

The bark beetle larvae spend their entire life gregariously feeding and developing under the bark. The larvae acquire microbial communities during feeding that mainly aid in nutrient acquisition, detoxification of plant defensive compounds (27, 66, 67) For instance, a significantly high abundance of *Pseudomonas*, *Pseudoxanthomonas*, *Endobacter* (Table 3) in pine-feeding *Ips* larvae documented in our study might be associated with lignocellulose and cellulose/hemicellulose degradation (21, 68, 69). In addition, *Pseudomonas* have also been reported to perform different ecological functions within bark beetle holobiont, including detoxification of tree chemical defences and protection against entomopathogenic fungi (70).

During metamorphosis, beetles undergo complete structural changes from larvae to adults via the non-feeding pupal stage, which might lead to the gain or loss of certain microbiota (71). The compartmentalisation of the internal structures and organogenesis during metamorphosis leads to different physiological conditions, including redox potential, oxygen concentrations, and pH changes influencing the distribution and survival of the insect microbiota (72). Recent findings by Peral-Aranega et al. (73) revealed that the bacterial diversity in *I. typographus* reduced in the pupal stage compared to the larvae and regained in the adult stages. However, our study observed no such trend in the two pine-feeding *Ips* bark beetles. It can be assumed that pine beetle larvae can accumulate bacterial partners during development and feeding under the bark, leading to high titers in pupae, as documented in a recent study (29). Furthermore, adult beetles perform multiple duties on maturation, including feeding, host finding, and reproduction. Such responsibilities of the adult beetles can be associated with the maintenance and selection of their symbionts (74). For instance, *Mycobacterium,* highly abundant in *I. acuminatus* adults, is known for its high adaptive competency (75); therefore, it can be assumed that it might help the beetle overcome unfavourable biotic and abiotic challenges during host-seeking. Interestingly, certain bacterial genera such as *Serratia*, *Pseudomonas*, *Dyella,* with differences in their relative abundance, are persistent across the life stages, suggesting their pivotal role in bark beetles holobiont (73). However, such possibilities need further experimental validation.

The bacterial richness and diversity varied across the same developmental stages between the two *Ips* beetles. For instance, our data revealed high bacterial richness in *I. sexdentatus* (ISX) larvae compared to *I. acuminatus* (IAC), while the bacterial diversity in IAC larvae was higher. This interspecies variation in the bacterial community might be associated with their variable preference for pine trees. Although both *I. sexdentatus* and *I. acuminatus* feed on the same pine trees, they have particular preferences for pine trees.

*I. acuminatus* beetles can attack healthy trees, whereas *I. sexdentatus* beetles are considered a secondary pest that prefers weakened and stressed pine trees to colonize (76, 77). In addition, *I. acuminatus* beetles infest young pine stands and plantations more aggressively than *I. sexdentatus* (14). Therefore, both *Ips* species might have differential resistance to host chemical defence that can influence bacterial partner selection and maintenance. Also, *I. acuminatus* beetles might require a wide range of bacterial species to deal with pine allelochemicals, which might explain the higher bacterial diversity in *I. acuminatus* larvae compared to *I. sexdentatus* larvae. In addition, *I. acuminatus* adult (wild) feeds on the upper canopy of the pine tree where the bark is thin, whereas *I. sexdentatus* adult (wild) beetles feed on the lower part of the bole where the bark is thick (14). Such habitat specialization can be reflected by the differences in the bacterial diversity and richness between the two bark beetle species feeding on the same host. Therefore, it can be assumed that microclimatic conditions can influence the bacterial richness and diversity in these beetles. However, such interpretations need to be further examined.

Furthermore, for bark beetles and many other species, the competency of an organism to associate with other microorganisms can allow the species to thrive under challenging scenarios like climatic fluctuations and resources (78). Therefore, the fungus-feeding nature of *I. acuminatus* larvae might have some influence on its bacterial population dynamics. The fungal species are pathogenic to the host tree, while on the contrary, they play nutritional roles in the *I. acuminatus* larvae (79) and might also interact with the bacterial communities residing in the developing larvae (80). This could explain the higher bacterial richness in fungus-feeding *I. acuminatus* larvae but a higher bacterial diversity in *I. sexdentatus* larvae. However, dedicated investigations are needed to confirm such possibilities experimentally *in beetlo*.

Laboratory adaptation and breeding conditions are essential factors that can affect bacterial associations unpredictably (29, 81). Compared to laboratory-bred beetles, broader and continuous environmental challenges in wild-collected beetles can encourage highly diversified and rich bacterial assemblage in wild-collected beetles. Furthermore, the bottleneck effect and high selective pressure are the key drivers in laboratory populations that often reduce symbiont load (81). Such observations may result in *I. sexdentatus* adults (wild) having higher bacterial diversity and richness than lab-bred adults (Figure 3A, Table 1, Figure 4C). However, no such trend was observed in *I. acuminatus*. Hence, dedicated investigations are needed to understand the ecological relevance of such observation. It is worth mentioning that the beetles were collected in different years; hence, the observed variation can also be due to feeding on different wood under different environmental conditions. A comparison of lab-bred (control wood, fed wood) and wild-collected (*I. sexdentatus* control and fed wood) wood samples for the same species (*I. sexdentatus*) has revealed that each wood sample poses a distinct bacterial population (Supplementary Figure 4, Supplementary excel 9) endorsing our prediction.

Although both pine beetles have distinct bacterial communities influenced by microclimatic conditions, canopy preference, metamorphosis, and feeding behaviour, our study revealed a core bacteriome across all developmental stages of two pine beetles (Figure 2C) that were presumed to perform conserved functions. Such observation was also documented as symbiotic partners to the other members of the same subfamily, endorsing their conserved functional roles in survival, growth, and development (82). For instance, core members like *Pseudomonas* and *Serratia* can degrade and utilize plant defence chemicals (monoterpenes) to produce anti-aggression pheromone-verbenone (17, 18, 83). Moreover, both bacterial genera can degrade complex polymers such as cellulose, starch, xylan, and lignin to help beetle nutrient acquisition (21, 65, 84). These bacteria genera can also recycle uric acid from beetle faeces, helping the beetles meet the crucial nitrogen requirement (23, 85) and producing antifungal compounds to suppress pathogenic fungal growth, aiding the beetle survival (86). Other core members such as *Dyella, Pseudoxanthomonas, Sphingomonas, Saccharomonospora, Acidisoma* have the enzymatic competency to help the beetle metabolize complex carbohydrates found in the conifer cell wall (21, 69, 87–90). Therefore, such microbes belonging to the core bacteriome can be further evaluated *in beetlo* for potential benefits and targeted for future microbes-mediated sustainable beetle management practices (91).

Our study also reveals beetle-mediated alterations in wood bacterial assemblage, similar to insect herbivores that reshape the native plant leaf microbiome (92). However, the degree of overlap or distinctiveness between insect and host (wood) microbiome remains ambiguous (93). Precisely, higher sharing between fed wood and beetle samples followed by higher bacterial diversity in fed wood samples than in control wood can be due to several reasons. It can be presumed that bacterial association in both beetle species is influenced by the horizontal transfer of bacteria from the pine host. Similarly, beetle-feeding introduces new members to the microhabitat (gallery wood) and vice versa. The abundance and richness of gallery wood bacteriome may be due to the beetle-introduced microbiome during feeding. Moreover, the beetle ecto-symbiont bacterial population can also have an influence. Further studies are required to comprehend the ecological relevance of beetle-mediated tree bacteriome alteration.

The abundance of specific bacterial ASVs involved in amino acid biosynthesis, cell wall degradation, and vitamin synthesis might endorse their role in nutritional supplementation (66, 85). Oxidoreductase enzymes that catalyze the transfer of electrons from one molecule to another were found to be the dominant enzymatic class (EC) between larvae and adult (p<0.001) stages, *I. acuminatus* larvae and pupae (p<0.001) (Figure 8, Supplementary excel 14,15). A higher abundance of Flavin-dependent oxidoreductase is documented in *I. acuminatus* larvae and pupae (Supplementary figure 6) that are reported to be crucial in several fundamental biological processes, including detoxification, catabolism, and biosynthesis (94, 95). Interestingly, the KEGG database predicted a higher abundance of oxidoreductase genes like OAR1 in *I. sexdentatus* adults. OAR1 in *Pseudomonas* genome are involved in fatty acid synthesis (96). However, in *I. sexdentatus* larvae, the dominating class was transferases (enzymatic class catalyzing the transfer of a group of atoms) (Supplementary excel 13, 15). The dominance of the transferases class in *I. sexdentatus* larvae can be reflected by a higher abundance of AspAT, a crucial enzyme in the Krebs cycle, and the synthesis of essential amino acids (97). Such observation in *I. sexdentatus* larvae might also be related to a high abundance of glycosyltransferase genes involved in cell wall biosynthesis during pine host feeding (Supplementary figure 6). The hydrolytic hexosaminidase gene was abundant in *I. sexdentatus* pupae (Supplementary figure 7) and is reportedly involved in bacterial chitin degradation (98). Concomitantly, between *I. sexdentatus* pupae and *I. acuminatus* pupae (p<0.001), *I. sexdentatus* adults (wild) and *I. acuminatus* adults (wild) (p<0.05) transferases and hydrolases were the dominant enzyme class. The dominance of transferase is consistent with the high abundance of N-acetyltransferase and glutathione S-transferase (GST) in the pupal stage of the two pine-feeding beetles (Supplementary figure 6). Acetyl-transferase enzymes help in several acetylation processes, including post-translational modification and multiple phosphorylation processes (99). Glutathione S-transferase is a vital cytosolic detoxification enzyme that can be functional against pine monoterpenes (100). Nevertheless, the EC data revealed that the enzymatic categorization of bacterial ASV abundance varied greatly and showed species and stage-specific abundance correlated with their physiological needs, such as monoterpene-rich host feeding. However, it is worth mentioning here that PICRUSt2 is a tool to predict the putative functional role of the associated bacterial communities based on marker genes. Further validation at the functional level based on culture-based techniques, metatranscriptomic and metaproteomic approaches is required.

### Conclusion

Our comparative metagenomic study reported life stage-specific and core bacterial genera and their putative ecological role in two pine-feeding bark beetle holobionts. It endorses the hypotheses that microclimatic conditions, life stages, and lab-breeding impact the pine beetle bacteriome. It also demonstrated that beetle feeding substantially influenced the host bacteriome at the feeding galleries. The metagenomic sequencing results were corroborated by quantitative PCR data, revealing a similar trend in the relative abundance of selected bacterial communities in the two pine-feeding beetles. Current data will facilitate further downstream studies (metatranscriptomics, metaproteomics, and culture-based bacterial functional characterization) to characterize the critical bacterial species aiding these *Ips* beetles thriving on defence-rich pine trees. Such essential bacterial partners can be used for microbiome-based sustainable pest management practices (91, 101, 102). Finally, such knowledge can serve as a primer to decipher the underlying microbial interactions at the tree-beetle interface and facilitate the integration of tree-beetle-microbiome interaction data into spatial models of forest ecosystem dynamics predictions.

## Data Availability

The bacteriome dataset in the study is available under NCBI Bio-project PRJNA854390.

## Acknowledgements

We appreciate the helpful feedback provided by the reviewers and handling editor. We also acknowledge Dr Jiri Synek, FLD, CZU, for beetle collection and maintenance assistance.

## Author Contributions

AR and AC conceptualization. AR, AK, RM sampling. AK experimental work. AR, AK and RM figure preparations and qPCR statistics. AR, AC and AK data interpretation and manuscript writing. All authors read and approved the final version of the manuscript.

## Publisher’s Note

All assertions in this article are exclusively those of the writers and do not necessarily reflect the views of their affiliated organizations, the publisher, editors, or reviewers. The publisher does not guarantee or support any product mentioned in this article or any claim made by its manufacturer.

## Abbreviation

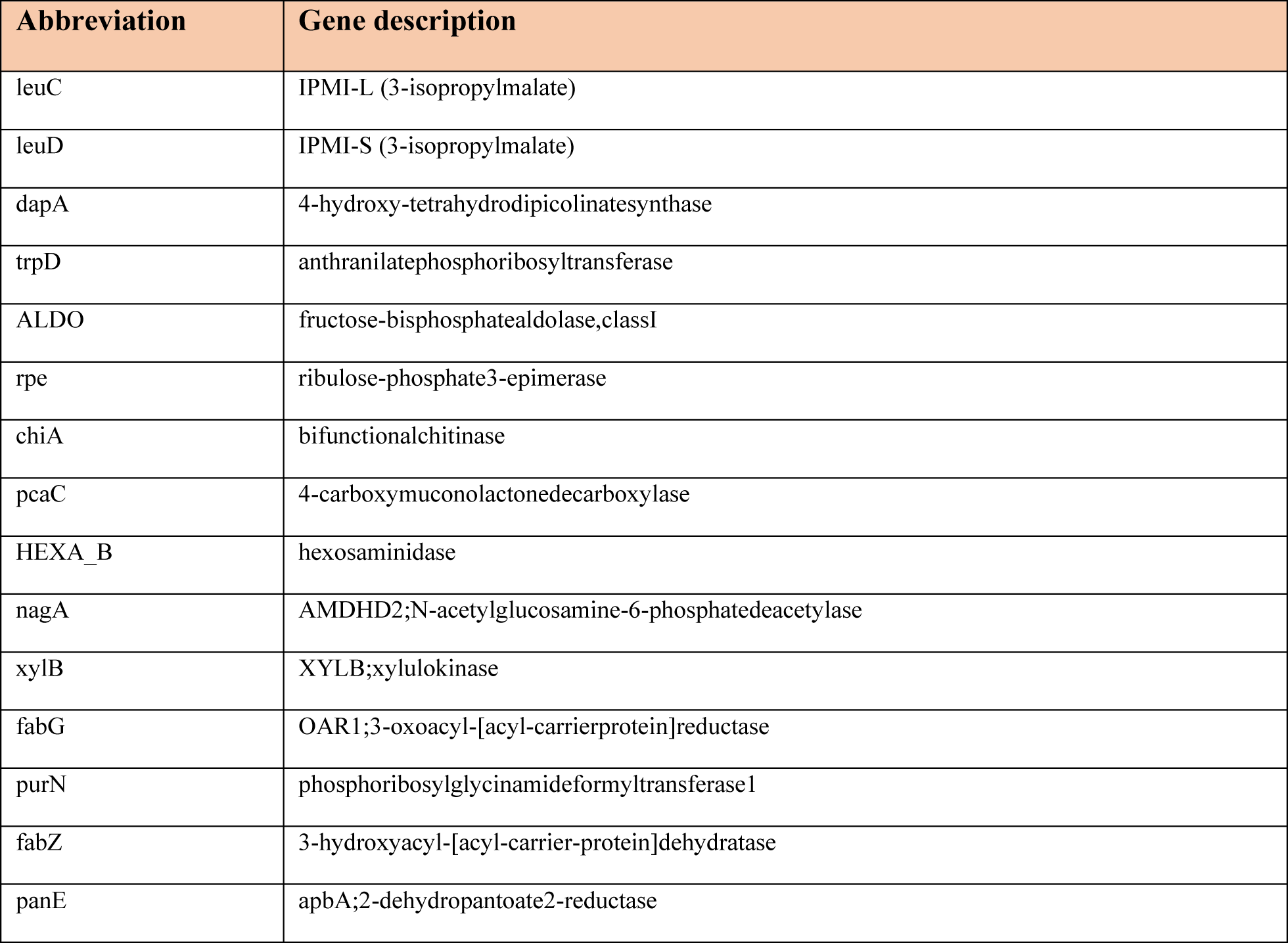

| Abbreviation | Gene description |
| --- | --- |
| leuC | IPMI-L (3-isopropylmalate) |
| leuD | IPMI-S (3-isopropylmalate) |
| dapA | 4-hydroxy-tetrahydrodipicolinatesynthase |
| trpD | anthranilatephosphoribosyltransferase |
| ALDO | fructose-bisphosphatealdolase,classI |
| rpe | ribulose-phosphate3-epimerase |
| chiA | bifunctionalchitinase |
| pcaC | 4-carboxymuconolactonedecarboxylase |
| HEXA_B | hexosaminidase |
| nagA | AMDHD2;N-acetylglucosamine-6-phosphatedeacetylase |
| xylB | XYLB;xylulokinase |
| fabG | OAR1;3-oxoacyl-[acyl-carrierprotein]reductase |
| purN | phosphoribosylglycinamideformyltransferaseI |
| fabZ | 3-hydroxyacyl-[acyl-carrier-protein]dehydratase |
| panE | apbA;2-dehydropantoate2-reductase |

## Supplementary figure legends

**Supplementary figure 1| Rarefaction curves.** Different samples are indicated by different colours and symbols.

**Supplementary figure 2| Boxplots depicting alpha diversity indices between different lab-bred, wild-type, wood tissue for two *Ips* pine beetles** (ISX-*Ips sexdentatus*, IAC-*Ips acuminatus*). Bacterial richness estimated by (A) Chao1 estimation indicating significant differences among different life stages, wild adults, and wood samples. Bacterial species evenness estimated by (B) Pielou estimation. The bacterial diversity is illustrated by (C) Shannon (D) Shannon index. Kruskal-Wallis-pairwise-group test was utilized to represent significant differences among different groups statistically. Different letters indicate statistical significance.

**Supplementary figure 3| Lefse analysis.** Histogram representing the LDA scores depicting the significantly abundant bacterial communities across (A) the different developmental stages of ISX (larvae, pupae, adult) (B) the different life stages of IAC (larvae, pupae, adult) (C) the lab-bred and wild adults of the two pine-feeding beetles, ISX and IAC (D) ISX wild adults along with the control wood and fed wood (E) IAC wild adults with the control wood and fed wood. The LDA score threshold value is set at log10>4, and the length of each bin illustrates the difference between the microbial biomarkers among the samples. (ISX-*I. sexdentatus*, IAC-*I. acuminatus*).

**Supplementary figure 4| Venn diagram illustrating the environmental impact (lab-breeding and wild collection) and beetle feeding on the wood bacterial communities** (ISX-*I. sexdentatus*).

**Supplementary figure 5| Estimation of overall bacterial abundance by quantitative PCR.** A) *I.sexdentaus* life stages; B) Wild vs lab-bred *I.sexdentaus* beetles; C) *I.sexdentaus vs I.accuminatus*.

**Supplementary figure 6| Functional prediction using PICRUSt2 based on (A) KEGG database and (B) COG database representing bacterial functional contribution in different beetle and wood samples.** (ISX-*Ips sexdentatus*, IAC-*I. acuminatus*).

**Supplementary figure 7| Significant abundance (p<0.05, t-test) of bacterial genes outlined from KEGG Enzyme database coding for enzymes involved in amino acid biosynthesis, cell wall degradation, vitamins and co-factors biosynthesis.** Heatmap represents the significant abundance of microbial genes among various life stages of ISX beetles and wild-type ISX. (ISX-*I. sexdentatus*)

**Supplementary figure 8| Significant abundance (p<0.05, t-test) of bacterial genes outlined from KEGG Enzyme database coding for enzymes involved in amino acid biosynthesis, cell wall degradation, vitamins and co-factors biosynthesis.** Heatmap depicting significant abundance of microbial genes among various life stages of IAC beetles and wild-type IAC. Darker colours represent higher abundance, whereas lighter colours indicate a lower abundance of a specific gene/enzyme. (IAC-*I. acuminatus*)

## Supplementary table legends

Supplementary Table 1|

**Bray-Curtis method was used for ADONIS analysis to evaluate the significant difference between different life stages of both *Ips* pine beetles** (ISX-*I. sexdentatus*, IAC-*I. acuminatus*) a**nd wood samples** (Df = degree of freedom, SS = sums of squares of deviations, MS = SS/Df, F. Model = F-test value, R2 = the ratio of grouping variance and total variance). Values in parentheses illustrate Residual Error. The p-value represents the significant variation between different stages of both bark beetles.

Supplementary Table 2|

**ANOSIM analysis represents the magnitude of variation between different life stages of both *Ips* pine beetles** (ISX-*I. sexdentatus*, IAC-*I. acuminatus*). In ANOSIM analysis, the positive R values indicate significant differences between the bacterial communities present at different life stages of both beetles. P-value < 0.05 indicates significant differences between different stages of both bark beetles and wood samples.

Supplementary Table 3| **Selected bacterial primers used for the quantitative PCR assay. Supplementary excel legends**

Supplementary excel 1| Raw and clean reads

Supplementary excel 2| ASV table indicating the relative abundance of bacterial communities in two pine-feeding beetles *I. sexdentatus* (ISX) and *I. acuminatus* (IAC), and the associated wood samples

Supplementary excel 3| Relative abundance of top 10 bacterial class

Supplementary excel 4| Core and Unique bacteriome in *I. sexdentatus* (ISX) and *I. acuminatus* (IAC) Supplementary excel 5| Alpha diversity indices

Supplementary excel 6 | Common and unique bacteriome among different stages of *I. sexdentatus* (ISX), *I. acuminatus* (IAC)

Supplementary excel 7 | | Core and unique bacterial communities illustrating the effect of lab breeding on the pine-feeding beetle bacteriome

Supplementary excel 8| | Common and unique bacteriome representing the host contribution to beetle bacteriome

Supplementary excel 9| Core and Unique bacterial communities indicating the environmental impact on wood microbiome

Supplementary excel 10| Statistical analysis on quantitative PCR data.

Supplementary excel 11| PICRUSt2 data representing the significantly abundant bacterial genes between different stages of *I. sexdentatus* (ISX)

Supplementary excel 12| PICRUSt2 illustrating the significantly abundant bacterial genes between different stages of *I. acuminatus* (IAC)

Supplementary excel 13| 4-fold change between different stages of *I. acuminatus* (IAC) EC abundance Supplementary excel 14| 4-fold change between different stages of *I. sexdentatus* (ISX) EC abundance

Supplementary excel 15| 4-fold change between different stages of *I. acuminatus* (IAC) and *I. sexdentatus* (ISX) EC abundance

